# LAP1 supports nuclear plasticity during constrained migration

**DOI:** 10.1101/2021.06.23.449503

**Authors:** Yaiza Jung-Garcia, Oscar Maiques, Irene Rodriguez-Hernandez, Bruce Fanshawe, Marie-Charlotte Domart, Matt Renshaw, Rosa M Marti, Xavier Matias-Guiu, Lucy M Collinson, Victoria Sanz-Moreno, Jeremy G Carlton

## Abstract

Metastasis involves dissemination of cancer cells away from a primary tumour and colonisation at distal sites. During this process, cancer cells must negotiate multiple physical constraints imposed by the microenvironment and tissue structure. The biophysical properties of the nucleus must be tuned since they pose a challenge to constrained migration. By analysing nuclear genes upregulated during the acquisition of metastatic potential, we discovered increased expression of the inner nuclear membrane protein LAP1 in metastatic melanoma cells and at the invasive fronts of human primary tumours and in metastases. Human cells express two LAP1 isoforms (LAP1B and LAP1C), which differ in their amino terminus. We found that whereas the longer isoform, LAP1B, binds more strongly to nuclear lamins and has restricted motility within the nuclear envelope, the shorter isoform, LAP1C, favours nuclear envelope blebbing and allows migration through constraints. We propose that LAP1 renders the nucleus plastic and enhances melanoma aggressiveness.

## INTRODUCTION

Metastatic spread accounts for the majority of cancer-related deaths ^1, 2^ and there is an urgent need to understand how metastatic potential is acquired. Metastatic melanoma is the leading cause of death for skin cancers ^3, 4^. Melanoma cells can switch between different collective and individual migratory modes, can degrade the extracellular matrix, and can reprogram cells in the tumour microenvironment to favour cancer cell survival, migration and invasion ^5-11^. Physical constraints, for example traversing tissue constrictions or the vascular endothelium, are a major barrier to metastatic spread ^12, 13^ and cells must negotiate multiple constraints before colonising new sites. While the cytoplasm can accommodate large deformations, the biophysical properties of the nucleus make translocation of this organelle the rate-limiting step during constrained migration ^13^.

Nuclear mechano-properties are regulated through protein-protein interactions between integral nuclear envelope (NE) proteins, the nuclear lamina and the cytoskeleton ^14-20^. Deficiencies in lamins can render NE membranes prone to rupture under mechanical stress, leaving genomic DNA exposed to damaging agents in the cytoplasm and prone to persistent damage due to the mis-localisation of repair factors ^21-25^. NE ruptures occur typically at NE blebs, where local remodelling of the lamina allows NE and chromatin herniation to protrude into the cytoplasm ^26^. As for plasma membrane blebs, actomyosin contractility regulates NE bleb dynamics ^26, 27^. However, whereas plasma membrane blebs can facilitate bleb-based migration ^11, 28-31^, whether NE blebs contribute to cellular migratory programmes is unclear. The involvement of nucleoskeletal and cytoskeletal rearrangements in NE bleb dynamics suggests that studying these structures might shed light into how tumour cells achieve a balance between nuclear stiffness and suppleness to allow constrained migration. Here, we challenged melanoma cells derived from primary tumours and those derived from isogenic metastatic lesions to multiple migratory constraints to identify proteins that enable cells to negotiate these challenges. Using a combination of transcriptomics, biochemistry, microscopy and live-cell imaging we discovered that the inner nuclear membrane (INM) protein lamin-associated polypeptide 1 (LAP1) enhances the ability of metastatic cells to undergo repeated constrained migration and tissue invasion through a migratory programme involving NE blebbing.

## RESULTS

### Metastatic melanoma cells negotiate repetitive constrained migration challenges

Using a pair of isogenic melanoma cell lines from the same patient (WM983A, derived from the primary tumour; WM983B, derived from a metastatic lesion) stably expressing a nucleus-localised GFP (GFP-NLS), we designed a multi-round transwell assay (Fig.1a) to understand the differential abilities of these cancer cells to negotiate repeated constraints during migration. Using transwells with a range of pore sizes smaller than the average nuclear diameter of tumour cells ^13^, we found that during the first round of migration, whilst decreasing pore size impaired migration, metastatic melanoma WM983B cells were more effective at negotiating constraints than primary melanoma WM983A cells (Fig.1b and Supplementary Fig.1a). Pore transit of WM983B cells was accompanied by enhanced NE blebbing (Fig.1c and Supplementary Fig.1b, c). On a second round of migration using sequentially 8-μm and 5-μm transwells, we found that while WM983B cells could negotiate this second challenge with a similar efficiency to the first challenge, WM983A cells were severely compromised by this second challenge (Fig.1d, e and Supplementary Fig.1d). In addition, pore transit of WM983B cells noticeably enhanced NE blebbing (Fig.1f and Supplementary Fig.1e, f) suggesting that NE blebbing either promotes the migratory ability of metastatic melanoma cells, or that negotiating the constraint induces NE blebbing.

**Figure 1.**
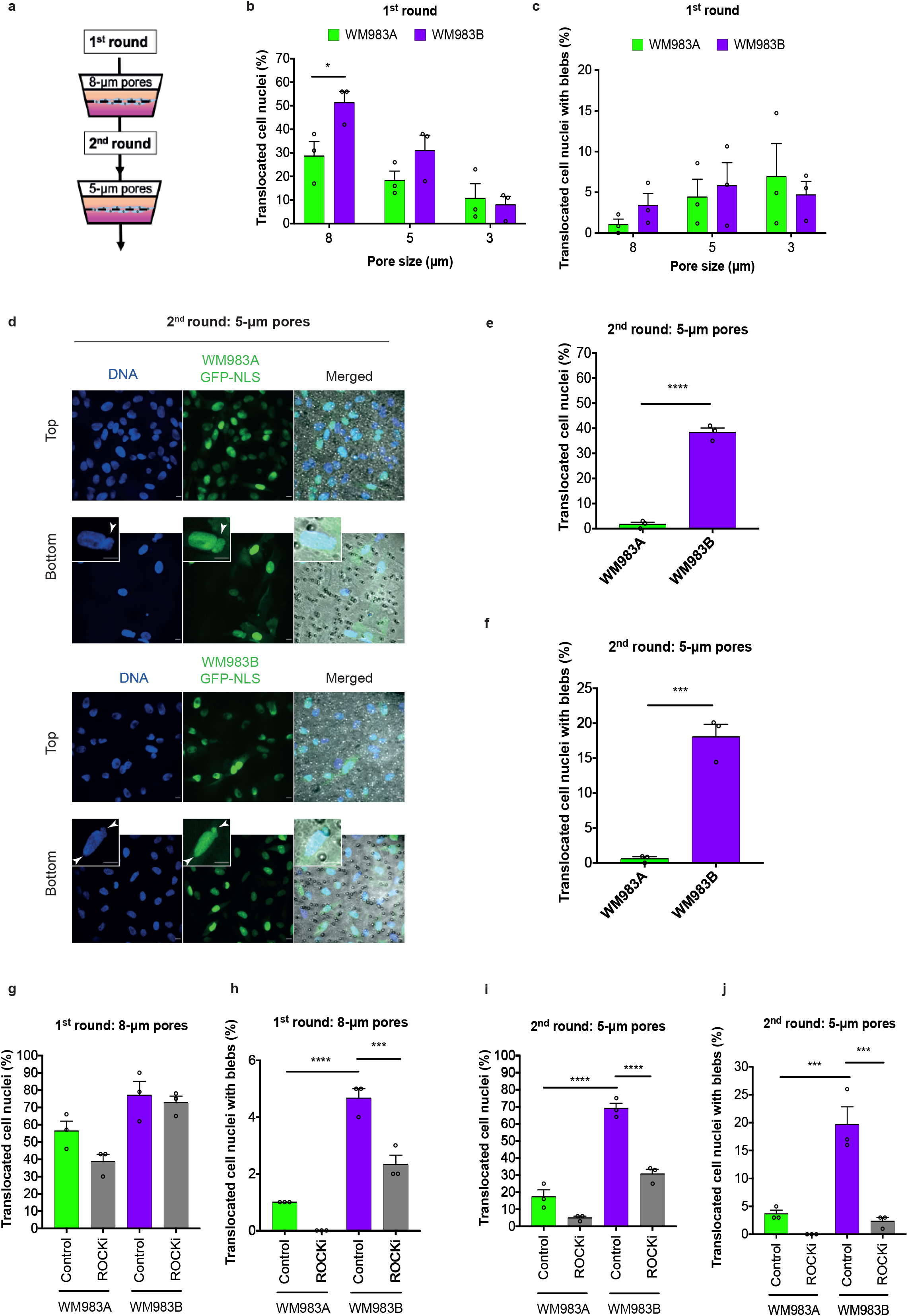
Metastatic melanoma cells can better negotiate repetitive constrained migration challenges than primary melanoma cells. **(a)** Schematic of transwell assays. Briefly, cells were challenged to migrate through transwells once or collected after a first round of migration and challenged repeatedly. The arrows indicate the direction for chemotactic migration. **(b)** Percentage of primary melanoma WM983A cells and metastatic melanoma WM983B cells that translocated their nuclei after a first round of migration in transwells of various pore sizes. **(c)** Percentage of WM983A and WM983B cells that translocated their nuclei and displayed nuclear envelope blebs after a first round of migration in transwells of various pore sizes. n= 1758 and 1728, respectively. **(d)** Representative pictures of WM983A and WM983B cells expressing GFP-NLS (green) and stained for DNA (blue) after a second round of transwell migration. Transwell pores could be visualised using transmitted light. The magnifications show nuclei that translocated and display nuclear envelope blebs indicated by white arrow heads. Scale bars, 30 μm and 10 μm magnifications. **(e)** Percentage of WM983A and WM983B cells that translocated their nuclei after a second round of transwell migration. **(f)** Percentage of WM983A and WM983B cells that translocated their nuclei and displayed nuclear envelope blebs after a second round of transwell migration. n= 192 and 313, respectively. **(g)** Percentage of WM983A and WM983B cells that translocated their nuclei after one round of transwell migration upon ROCK inhibitor (ROCKi) GSK269962A treatment. **(h)** Percentage of WM983A and WM983B cells that translocated their nuclei and displayed nuclear envelope blebs after one round of transwell migration upon ROCKi treatment. n= 1689 and 1757, respectively. **(i)** Percentage of WM983A and WM983B cells that translocated their nuclei after a second round of transwell migration upon ROCKi treatment. **(j)** Percentage of WM983A and WM983B cells that translocated their nuclei and displayed nuclear envelope blebs after a second round of transwell migration upon ROCKi treatment. n= 1119 and 1487. Experimental data have been pooled from three individual experiments. Graphs show the mean and error bars represent SEM. p values calculated by one-way ANOVA, two-way ANOVA, and unpaired t test; *p < 0.05, ***p<0.001, ****p < 0.0001.

We confirmed that WM983A cells and WM983B cells retained this migratory behaviour after a third round of constrained migration using sequentially 8-μm, 8-μm and 5-μm pores (Supplementary Fig.1g, h). The percentage of translocated nuclei exhibiting NE blebs after this third transit was again enhanced relative to the cells that did not translocate their nuclei, but was not enriched, suggesting that repeated constrictions are not selecting for a population of cells with elevated NE blebs (Supplementary Fig.1i, j). We confirmed that both cell lines have a similar number of viable cells after migration (Supplementary Fig.2a, b). We also confirmed that neither the morphological features of apoptosis nor active Caspase 3 were present in either of the melanoma cell lines, before or after a second round of migration (Supplementary Fig.2b, c), suggesting that passage through repeated constraints does not activate a cell death programme.

WM983B cells display enhanced Rho-ROCK1/2-driven Myosin II (MLC2) activity compared to WM983A cells ^8, 9, 32^ (Supplementary Fig.2d). We reasoned that higher MLC2 activity hence higher actomyosin contractility could contribute to both the generation of NE blebs and the enhanced migration of WM983B cells over WM983A cells. To test this hypothesis, we treated cells with the ROCK1/2 inhibitor (ROCKi) GSK269962A and challenged them sequentially to migration through 8-μm and 5-μm transwell pores. We confirmed that MLC2 activity was reduced after ROCK1/2 inhibition (Supplementary Fig.2d, e) and found that ROCK1/2 inhibition did not reduce nuclear translocation but did reduce NE blebbing of WM983B cells during the first round of migration (Fig.1g, h). However, ROCK1/2 inhibition markedly impaired nuclear translocation and reduced NE blebbing after pore transit during the second round (Fig.1i, j), suggesting that passage through the first constraint activates a Rho-ROCK1/2-dependent migration programme for subsequent passages. We concluded that Rho-ROCK1/2-driven actomyosin contractility is required for NE bleb generation and nuclear translocation of melanoma cells through repeated constraints.

### The nuclear envelope of metastatic melanoma cells is highly dynamic

We next hypothesised that melanoma cells that had previously negotiated multiple constraints during metastasis *in vivo* may have retained a nuclear mechanical memory. We examined the nucleus of unconfined melanoma cells with variable metastatic potential to determine whether there were underlying differences in the degree of NE blebbing. Lamin A/C immunofluorescence revealed that 30% of WM983B cells display NE blebs compared to only 5% of WM983A cells (Fig.2a). We defined four main NE bleb categories according to bleb shape (Fig.2b, c) and discovered that there were more NE blebs per nucleus and higher bleb phenotypical variability in WM983B cells compared to WM983A cells (Fig.2c) re-enforcing the association of NE pleiomorphisms with melanoma aggressiveness. Furthermore, we scrutinised the NE of melanocytes and found NE blebs only in 1% of cells (Supplementary Fig.3a, b) suggesting that NE blebbing is a feature of malignancy.

**Figure 2.**
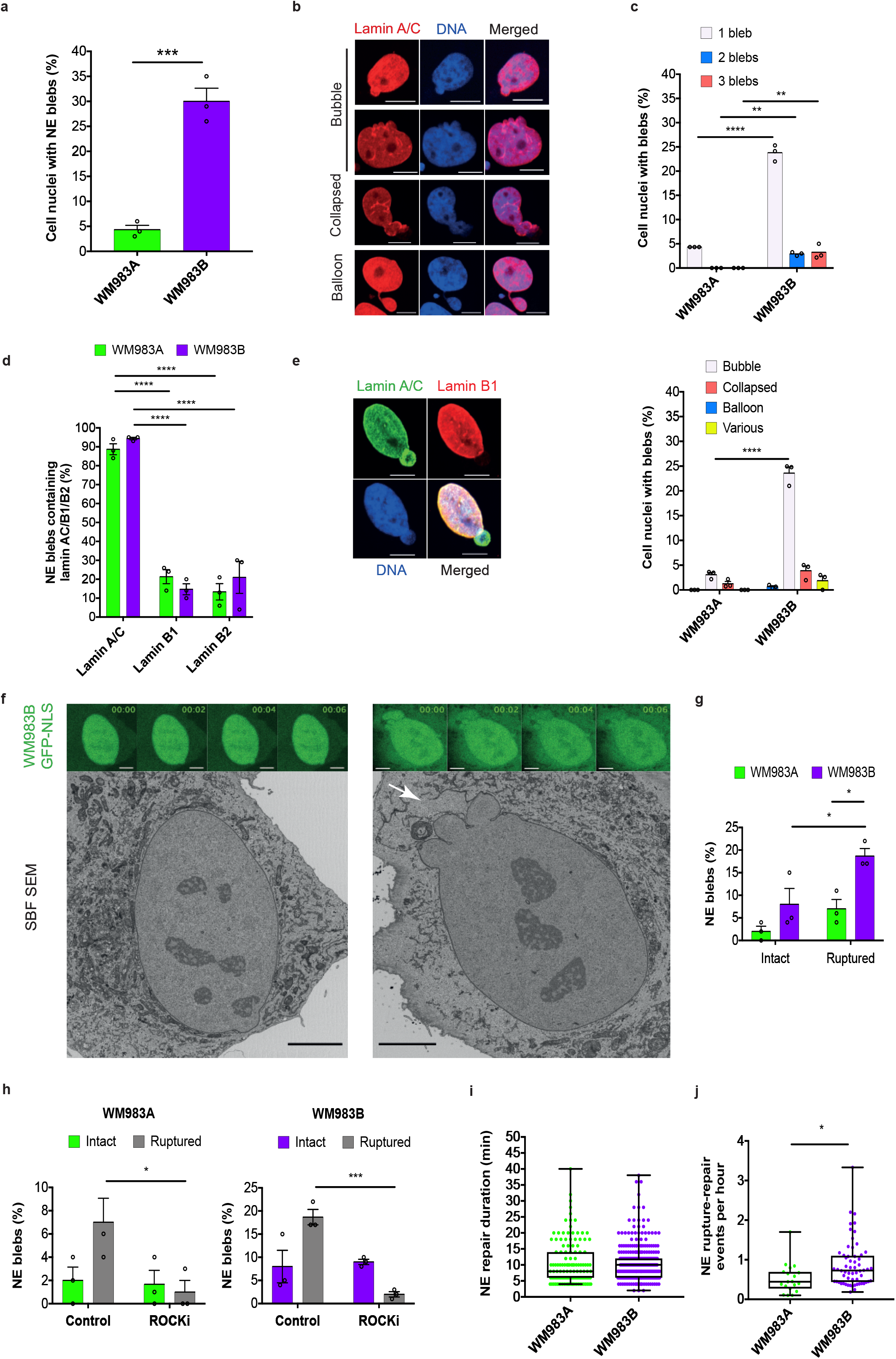
The nuclear envelope of metastatic melanoma cells is highly dynamic. **(a)** Percentage of primary melanoma WM983A and metastatic melanoma WM983B cells with nuclear envelope blebs. **(b)** Representative pictures of WM983B nuclei with nuclear envelope blebs of different shapes stained for Lamin A/C (red) and DNA (blue). Scale bars, 10 μm. **(c)** Percentage of nuclei with nuclear envelope blebs according to bleb number and bleb shape per nucleus in WM983A and WM983B cells. n= 434 and 561, respectively. **(d**) Percentage of nuclear envelope blebs containing Lamin A/C, Lamin B1 or Lamin B2 in WM983A and WM983B cells. n= 605 and 510, respectively. **(e)** Representative picture of a WM983B nucleus with a typical nuclear envelope bleb stained for Lamin A/C (green), Lamin B1 (red) and DNA (blue). Scale bars, 10 μm. **f)** CLEM of representative image sequences of WM983B GFP-NLS nuclei with an intact nuclear envelope bleb (left) and ruptured nuclear envelope blebs (right) and representative SBF SEM images of the same nuclei. The white arrow indicates the site of rupture of the nuclear membranes. Scale bars, 5 μm. **(g)** Percentage of intact and ruptured nuclear envelope blebs in WM983A and WM983B cells over the course of 15 hours. **(h)** Percentage of intact and ruptured nuclear envelope blebs in WM983A and WM983B after treatment with ROCK inhibitor (ROCKi) GSK269962A over the course of 15 hours. **(i)** Duration of nuclear envelope repair in WM983A and WM983B cells. **(j)** Nuclear envelope rupture-repair events per hour in WM983A and WM983B cells over the course of 15 hours. n= 1013 and 1008, respectively. Experimental data have been pooled from three individual experiments. **a, c, d, g, h** Graphs show the mean and error bars represent SEM. **i, j** Horizontal lines show the median and whiskers show minimum and maximum range of values. p values calculated by one-way ANOVA, two-way ANOVA, and unpaired t test; *p < 0.05, **p < 0.01, ***p < 0.001, ****p < 0.0001.

To assess NE blebbing in a more physiological setting, we examined melanoma cells seeded in collagen-I matrices. We confirmed that WM983B cells in collagen I were more amoeboid and harboured higher MLC2 phosphorylation levels than WM983A cells (Supplementary Fig.3c-e, i). WM983B cells in collagen had a more irregular nucleus, with 20% displaying NE blebs compared to 10% of WM983A cells and a higher karyoplasmic ratio (Supplementary Fig.3j-l, p). We obtained similar results with another melanoma cell line pair (highly metastatic and highly amoeboid A375M2 cells and less metastatic and less amoeboid A375P cells ^9, 32, 33^ (Supplementary Fig.3f-I, m-p)). Therefore, we suggest that the more metastatic and amoeboid the melanoma cells are, the more likely they are to display NE blebs.

NE blebs form at regions of the NE with compromised structural integrity ^21-24, 26^. We investigated if NE blebs in melanoma cells were associated with local changes in nuclear lamins. We found that about 90% of NE blebs present a weak or absent B-type lamin staining but maintain persistent Lamin A/C staining (Fig.2d, e). In addition, chromatin extruded in NE blebs and was frequently positive for markers of double-strand DNA breaks (Supplementary Fig.4a, b), which suggests that they are spots for local DNA damage generation. Nevertheless, overall levels of DNA damage were not increased (Supplementary Fig.4c).

We examined the dynamics of NE bleb formation, rupture and repair using unconfined melanoma cells stably expressing GFP-NLS. We found that NE blebs could either be intact or ruptured and this was further assessed using correlative light and electron microscopy (CLEM) demonstrating clear rupture of the NE at the bleb with concomitant leakage of the GFP-NLS signal (Fig.2f and Supplementary Movies 1a, 1b, 2a, 2b). We found that ruptured NE blebs were more frequent than intact blebs and that WM983B cells exhibited more ruptured NE blebs than WM983A cells (Fig.2g). We next tested if actomyosin-driven forces imposed on the nucleus were responsible for inducing NE blebbing and rupture. Indeed, both NE blebbing and ruptures were abrogated by ROCK inhibition (Fig.2h). To preserve cellular viability during interphase NE ruptures, cells reseal their NE via ESCRT-III-dependent repair ^24, 26^. We found that NE ruptures in melanoma cells are transient, and the average repair time was 10 minutes in both WM983A and WM983B cells (Fig.2i). Consistent with a higher background level of NE instability in WM983B cells, the rupture-repair rate for these cells was higher than in WM983A cells, with up to one event per hour (Fig.2j, Supplementary Fig.4d, e and Supplementary Movies 3, 4). These results suggest that NE ruptures occur at sites deficient in B-type lamins, that they are more frequent in metastatic cells and that an efficient NE repair programme acts to preserve cellular viability.

### Transcriptomics reveals *TOR1AIP1* upregulation in metastatic cells

As the degree of NE blebbing and migratory ability correlates with disease progression (Fig.1 and Supplementary Fig.1, 3), we interrogated a previously published transcriptional signature. Transcriptomes of amoeboid A375M2 melanoma cells were compared to intrinsically less amoeboid and less metastatic A375P melanoma cells with lower MLC2 activity ^6-8, 32^ using Gene Set Enrichment Analysis (GSEA) and focusing on genes encoding nuclear proteins (Fig.3a). This analysis showed that 63% of the gene sets containing genes encoding nuclear proteins were upregulated in A375M2 cells (FDR < 5%), when compared with A375P cells. Only 1% of these gene sets were upregulated in A375P cells compared to A375M2 cells (Fig.3b). Strikingly, about one third of upregulated gene sets in A375M2 cells were related to the nuclear membrane and organelle organisation (Fig.3b and Supplementary Tables 1-12). Leading-Edge Analysis ^34^ allowed us to identify a cluster of concurrently upregulated genes across NE gene sets (Fig.3c). Individual gene upregulation and statistical significance were evaluated and a final list of 7 candidate genes showing statistically significant upregulation was compiled (Fig.3e). Next, transcriptomes of A375M2 cells were compared to A375M2 cells treated with several contractility inhibitors (ROCK1/2 inhibitors H1152 or Y27632, or myosin II inhibitor blebbistatin), which revealed similar gene expression changes (Fig.3d, e). These data point at a specific transcriptional programme of genes encoding for nuclear proteins that is linked to high levels of actomyosin contractility. Candidate gene upregulation in A375M2 cells compared to A375P cells was confirmed by qPCR. Whilst all the genes were upregulated in A375M2 cells, *OSBPL8, SUMO1* and *TOR1AIP1* achieved statistical significance (Fig.3f). Exploring this in an orthogonal system, we confirmed statistically significant upregulation of *TOR1AIP1* by qPCR in WM983B cells compared to WM983A cells (Fig.3g).

**Figure 3.**
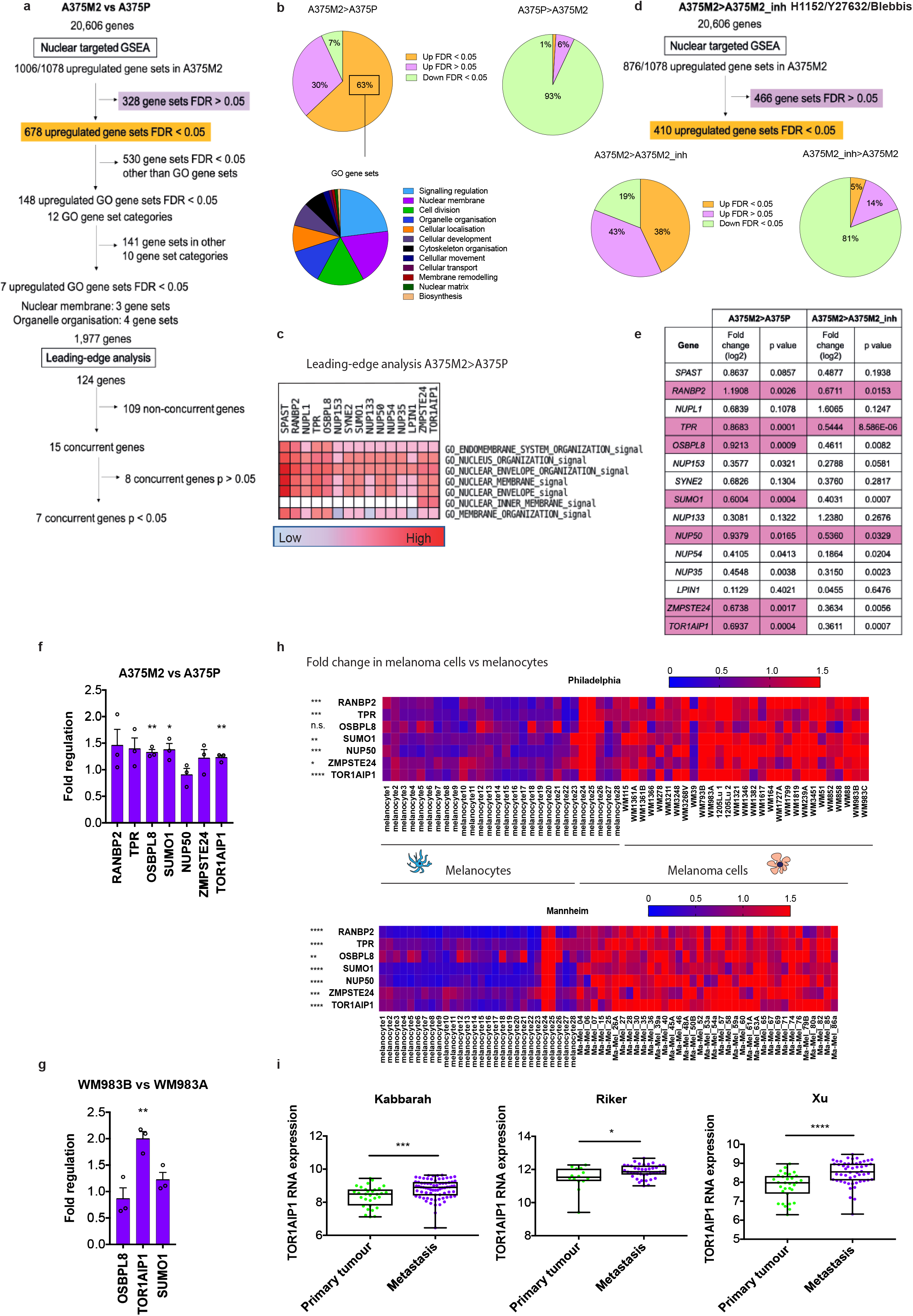
Nuclear transcriptomics reveals TOR1AIP1 upregulation in metastasis. **(a)** Flow diagram of GSEA carried out comparing the nuclear transcriptomes of highly amoeboid and highly metastatic melanoma A375M2 cells with less amoeboid and less metastatic melanoma A375P cells from data of gene expression microarray analysis. **(b)** The upper graphs show the distribution of upregulated and downregulated nuclear gene sets in A375M2 cells (left) and A375P cells (right). The lower graph shows the 12 most enriched GO gene set categories in A375M2. **(c)** Section of heatmap of the leading-edge analysis showing cluster of upregulated nuclear envelope genes in A375M2 cells. **(d)** Flow diagram of the GSEA carried out comparing A375M2 cells with A375M2 cells treated with contractility inhibitors (ROCK inhibitors H1152 or Y27632 or myosin inhibitor blebbistatin) (A375M2_inh). The graphs show the distribution of upregulated and downregulated nuclear gene sets in A375M2 cells (left) and A375M2_inh (right). **(e)** Upregulated nuclear envelope genes selected from leading-edge analysis with their log2 fold change in expression and statistical significance in A375M2 compared to A375P cells and in A375M2 compared to A375M2_inh. Statistically significant upregulated genes are coloured in pink in the table. **(f)** Fold regulation of candidate genes expression validated by qPCR in A375M2 compared to A375P. **(g)** Fold regulation of *OSBPL8, TOR1AIP1* and *SUMO1* expression validated by qPCR in metastatic melanoma WM983B cells compared to primary melanoma WM983A cells. **(h)** Heatmaps displaying fold change in expression of candidate genes in melanoma cell lines compared to melanocytes from Philadelphia and Mannheim datasets. **(i)** *TOR1AIP1* expression in primary tumours and metastasis in Kabbarah (n= 31 and 73, respectively), Riker (n= 14 and 40, respectively) and Xu (n= 31 and 52, respectively) melanoma patient datasets. Experimental data have been pooled from three individual experiments. **f, g** Graphs show the mean and error bars represent SEM. **i** Horizontal lines show the median and whiskers show minimum and maximum range of values. p values calculated by unpaired t tests; *p < 0.05, **p < 0.01, ***p < 0.001, ****p < 0.0001.

To assess how the expression of our candidate genes changes in melanoma progression, gene expression was analysed in two publicly available datasets (Philadelphia and Mannheim ^35^) containing transcriptomic profiles of melanoma cell lines compared to melanocytes. In both cases, upregulation of all our candidate genes was observed in melanoma cell lines (Fig.3h). Furthermore, using publicly available patient datasets (Kabbarah ^36^, Riker ^37^ and Xu ^38^), we found that *TOR1AIP1* mRNA levels were consistently upregulated in human samples obtained from metastatic melanoma lesions compared to primary melanoma (Fig.3i). These data suggest that expression of *TOR1AIP1* is upregulated in metastatic melanoma.

### LAP1 is overexpressed in melanoma metastasis

We next assessed the expression of the protein encoded by *TOR1AIP1*, LAP1, in human melanoma tumours grown in severe combined immunodeficient (SCID) mice. After subcutaneous injection of WM983A cells or WM983B cells, sections from such tumours were stained with antisera raised against LAP1. High-resolution immunohistochemistry coupled to digital pathology revealed that LAP1 localised to the NE of melanoma cells in these tumours. We observed that LAP1 expression was higher at the invasive fronts (IF) compared to the tumour bodies (TB) of these tumours (Fig.4a, b). We scored LAP1 intensity from 0 (very low) to 3 (very high) in individual tumour cell nuclei and observed that tumour cells showing the highest levels of LAP1 had a higher karyoplasmic ratio and represented a higher proportion in WM983B-derived tumours compared to WM983A tumours (Fig.4c and Supplementary Fig.5a). We confirmed these observations comparing A375M2-with A375P-derived tumours and found that they retained the same LAP1 staining pattern (Fig.4d-f and Supplementary Fig.5b).

**Figure 4.**
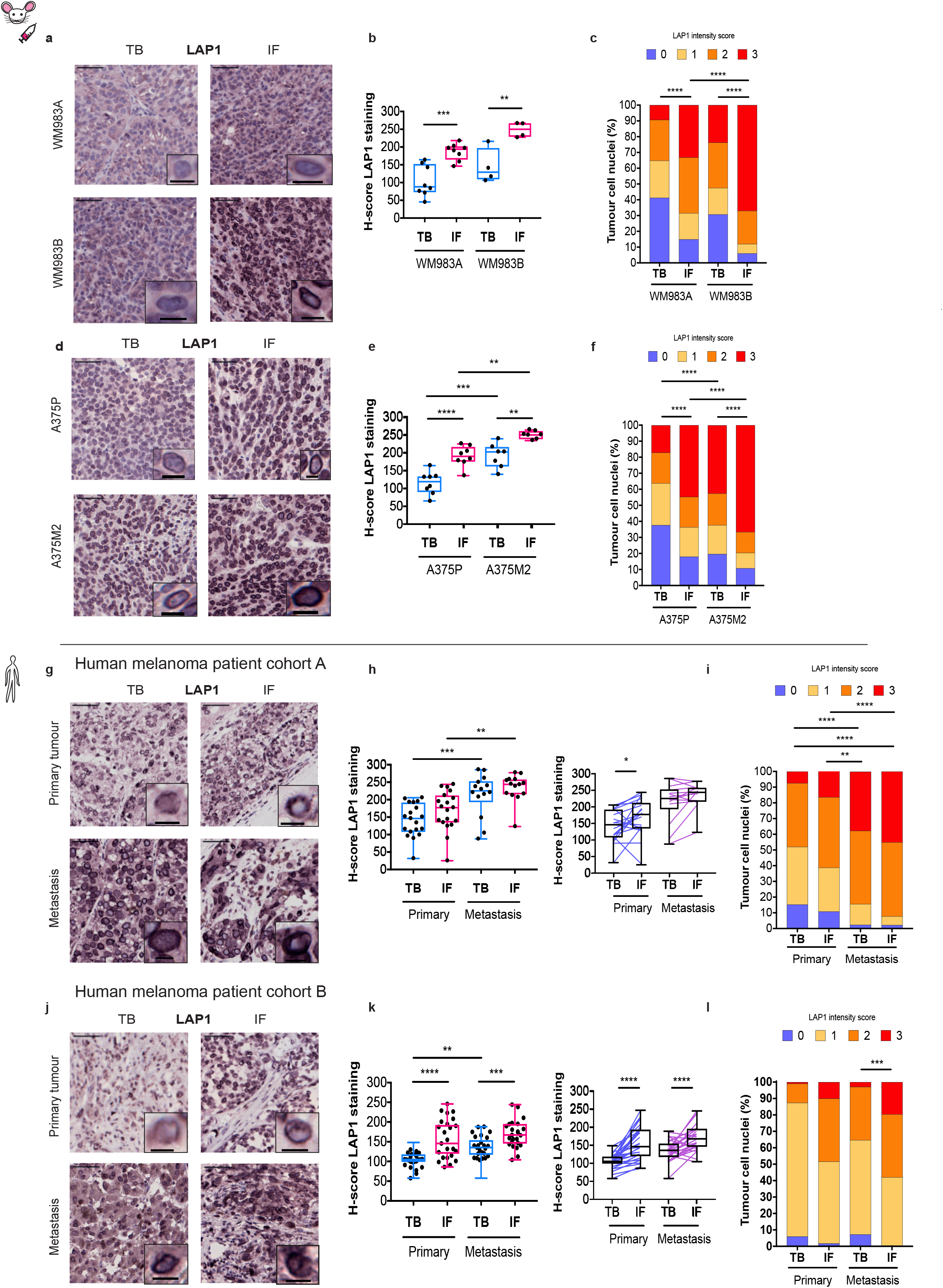
LAP1 is overexpressed in melanoma metastasis. **(a)** Representative images of tumour bodies (TB) and invasive fronts (IF) of WM983A and WM983B melanoma tumours grown in SCID mice. Scale bars, 50 μm. The magnifications show representative cell nuclei. Scale bars, 10 μm. **(b)** H-score for LAP1 staining in TBs and IFs of WM983A and WM983B tumours. **(c)** Percentage of tumour cell nuclei according to LAP1 intensity score in TBs and IFs of WM983A and WM983B tumours. n= 8 and 4, respectively. **(d)** Representative images of TBs and IFs of A375P and A375M2 melanoma tumours grown in SCID mice. Scale bars, 50 μm. The magnifications show representative cell nuclei. Scale bars, 10 μm. **(e)** H-score for LAP1 staining in TBs and IFs of A375P and A375M2 tumours. **(f)** Percentage of tumour cell nuclei according to LAP1 intensity score in TBs and IFs of A375P and A375M2 tumours. n= 8 and 7, respectively. **(g)** Representative images of TB and IF of a primary tumour and a metastasis in human melanoma patient cohort A. Scale bars, 50 μm. The magnifications show representative cell nuclei. Scale bars, 10 μm. **(h)** H-score for LAP1 staining in TBs and IFs of primary tumours and metastases in cohort A by unpaired (left) or paired (right) analysis. **(i)** Percentage of tumour cell nuclei according to LAP1 intensity score in TBs and IFs in cohort A. n= 19 primary tumours and 14 metastases. **(j)** Representative images of TB and IF of a primary tumour and a metastasis in human melanoma patient cohort B. Scale bars, 50 μm. The magnifications show representative cell nuclei. Scale bars, 10 μm. **(k)** H-score for LAP1 staining in TBs and IFs of primary tumours and metastases in cohort B by unpaired (left) or paired (right) analysis. **(l)** Percentage of tumour cell nuclei according to LAP1 intensity score in TBs and IFs in cohort B. n= 29 primary tumours and 29 metastases. **b, e, h, k** Horizontal lines show the median and whiskers show minimum and maximum range of values. p values calculated by one-way ANOVA, two-way ANOVA, and paired t-test; *p < 0.05, **p < 0.01, ***p < 0.001, ****p < 0.0001.

We next assessed LAP1 expression in human tissue from normal skin and melanoma. In agreement with our *in-silico* data for melanocytes, we found that LAP1 expression was very low in melanocytes compared to keratinocytes (Supplementary Fig.5c, d). We next used tissue microarrays from two human melanoma patient cohorts (cohort A including 19 primary tumours and 14 metastases and cohort B with a total of 29 primary tumours and their matched metastases ^32^) (Supplementary Tables 13, 14). We found increased LAP1 expression in IFs compared to the TBs of primary tumours and in metastatic lesions compared to primary tumours (Fig.4g-l). Importantly, we observed that tumour cells showing very high levels of LAP1 had a higher karyoplasmic ratio (Supplementary Fig.5e, f) and such cells were enriched in melanoma metastases compared to primary melanomas (Fig.4i, l).

### LAP1 is required for repeated constrained migration

Human cells express two isoforms of LAP1 that differ in the length of their amino terminus (NT); a long isoform, LAP1B, and a shorter isoform, LAP1C, generated by use of an alternative translation initiation at codon 122 ^39, 40^ (Fig.5a). We quantified the expression of LAP1B and LAP1C in melanocytes, primary melanoma cells and metastatic melanoma cells. Consistent with our transcriptomic analyses, we found that both LAP1 isoforms were upregulated in metastatic melanoma cells (Fig.5b, c).

**Figure 5.**
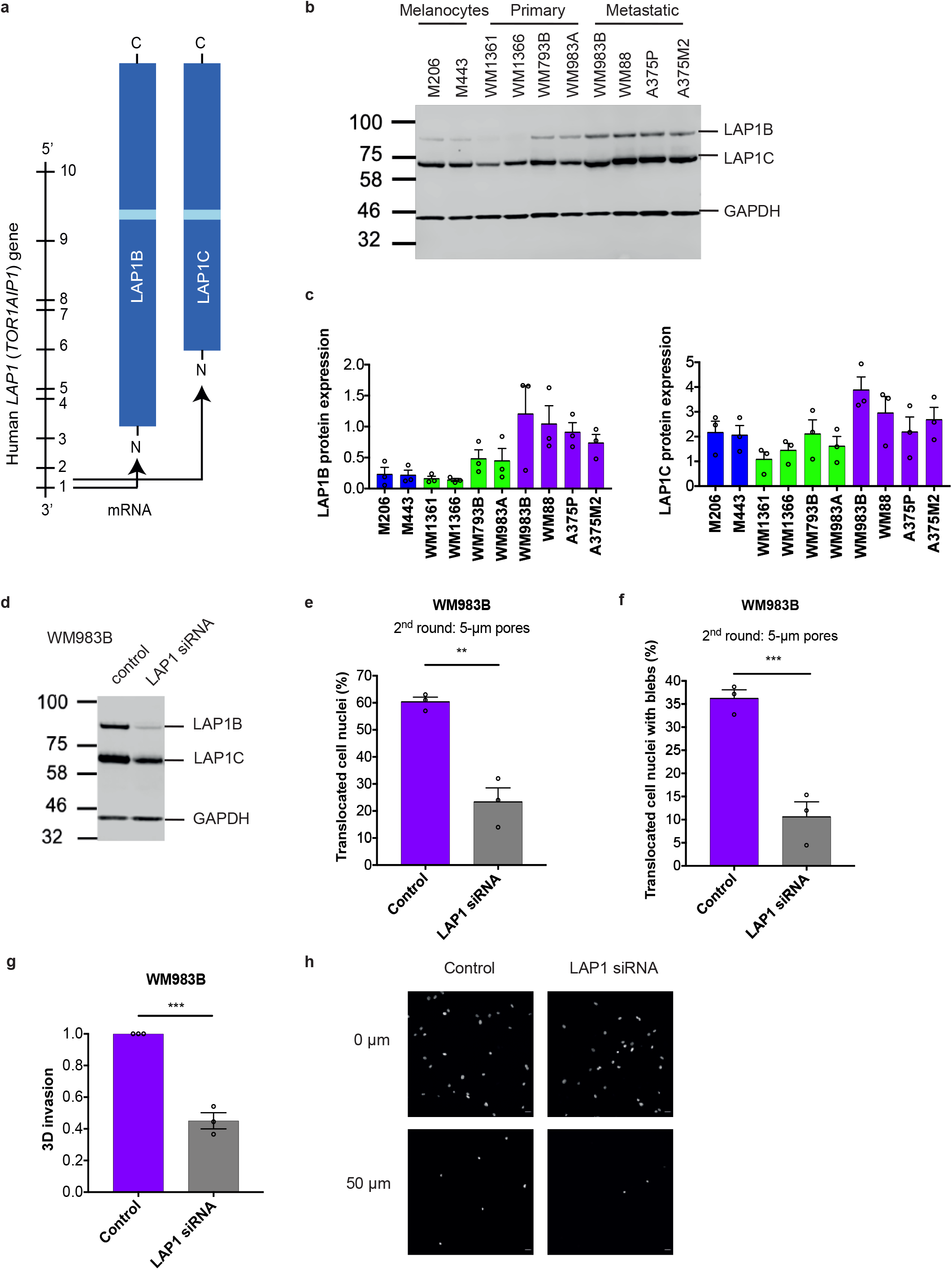
LAP1 is required for repeated constrained migration. **(a)** Schematic of *TOR1AIP1* human gene transcription and translation. The gene is comprised of 10 exons and encodes a long protein isoform (584 aminoacids), LAP1B, and a short protein isoform (462 aminoacids), LAP1C. There are two transcripts produced by alternative splicing and each gives rise to a LAP1B variant: a full-length variant, variant 1, and a variant missing one alanine at position 185, variant 2. Note that here only one transcript has been represented for ease of presentation. LAP1C results from an alternative translation initiation site at position 122. The transmembrane domain has been coloured in light blue. Proteins have been drawn at aminoacid scale. **(b)** Representative immunoblot for LAP1 expression levels in melanocytes, primary melanoma cells and metastatic melanoma cells. **(c)** Quantification of LAP1B and LAP1C protein expression in panel of cell lines in (b). **(d)** Representative immunoblot for LAP1 expression levels in WM983B cells upon LAP1 depletion with siRNA pool. **(e)** Percentage of WM983B cells that translocated their nuclei after a second round of transwell migration upon LAP1 depletion with siRNA pool. **(f)** Percentage of WM983B cells that translocated their nuclei and displayed nuclear envelope blebs after a second round of transwell migration upon LAP1 depletion with siRNA pool. n= 872 and 705, respectively. **(g)** Invasion index of WM983B cells in 3D collagen I matrices upon LAP1 depletion with siRNA pool. **(h)** Representative images of WM983B cells stained for DNA at 0 μm and up to 50 μm into collagen upon LAP1 depletion with siRNA pool. Scale bars, 30 μm. n= 673 and 539, respectively. Experimental data have been pooled from three individual experiments. Graphs show the mean and error bars represent SEM. p values calculated by one-way ANOVA and unpaired t test; **p < 0.01, ***p < 0.001.

We took a loss of function approach to understand whether LAP1 contributes to NE blebbing and the enhanced migration of WM983B cells. Whilst LAP1 isoforms were relatively resistant to siRNA depletion, we were able to reduce expression of LAP1 isoforms in WM983B cells to match the levels observed in WM983A cells (Fig.5d and Supplementary Fig.6a). We performed two-round transwell migration assays and discovered that reducing LAP1 expression levels in WM983B cells resulted in a reduction of 40% in second-round migration efficiency (Fig.5e) with no effect on cell viability (Supplementary Fig.6b). LAP1 depletion also supressed NE blebbing associated with the second round of transwell migration in WM983B cells (Fig.5f). We confirmed these results using two independent LAP1 siRNAs (Supplementary Fig.6c-f). We next performed 3D-invasion assays to assess the impact of LAP1 depletion on the ability of WM983B cells to invade through complex collagen-I matrices. We found that reduced LAP1 expression levels decreased the ability of WM983B cells to invade into 3D collagen I (Fig.5g, h). Overall, these data suggest that as well as being required for NE-bleb generation, LAP1 has a role in supporting the ability of metastatic melanoma cells to negotiate constraints in the microenvironment.

### LAP1B and LAP1C are differentially tethered to nuclear lamins

We next set out to understand if LAP1 isoforms play different roles based on the distinct length of their NTs. We carried out a solubilisation assay, and in agreement with previous studies ^39, 41, 42^ found that LAP1C was released from the nucleus of melanoma cells under mild extraction conditions, whereas LAP1B was only released when the nuclear lamina was solubilised (Supplementary Fig.7a). These data suggested that LAP1B might be involved in stronger protein-protein interactions with the nuclear lamina. The NT of LAP1 interacts with Lamin A/C and Lamin B1 ^41^ and two distinct lamin-binding regions in LAP1’s NT have been described: 1-72 (present in the unique region of LAP1B) and 184-337 (present in both LAP1B and LAP1C) ^43^. We hypothesised that the different length of the NTs of LAP1 isoforms might encode differential lamin-binding properties.

Solubilising INM proteins for classical immunoprecipitation approaches while retaining native interactions is challenging. We instead employed a mitochondrial retargeting assay and expressed GFP-Lamin A/C or GFP-Lamin B1 and either LAP1B^NT^, LAP1C^NT^ or the unique region of LAP1B’s NT (LAP1B^uNT^) fused to HA and the mitochondrial targeting sequence from Monoamine Oxygenase ^44^ (HA-MTS). We looked for mitochondrial re-localisation of GFP-Lamins or relocalisation of HA-tagged mitochondria to the nucleus and NE as a readout of this interaction (Fig.6a). We found that mitochondria presenting HA-LAP1B^NT^ or HA-LAP1B^uNT^ on their surface could be strongly relocalised to the GFP-Lamin positive NE, whereas mitochondria expressing HA-LAP1C^NT^ were less able to be relocated (Fig.6b, c). Consistent with the presence of a chromatin-binding region (CBR) in LAP1B’s NT ^43^, mitochondria presenting this region were strongly relocalised to the chromatin periphery. In all cases, expression of the HA-LAP1 NTs on mitochondria induced their clustering, but mitochondria that did not relocalise to the NE were able to strongly recruit GFP-Lamin B1, but not GFP-Lamin A/C. We suggest that as well as containing a CBR ^43^, the unique NT of LAP1B encodes a dominant lamin-binding domain that displays preference for B-type lamins.

**Figure 6.**
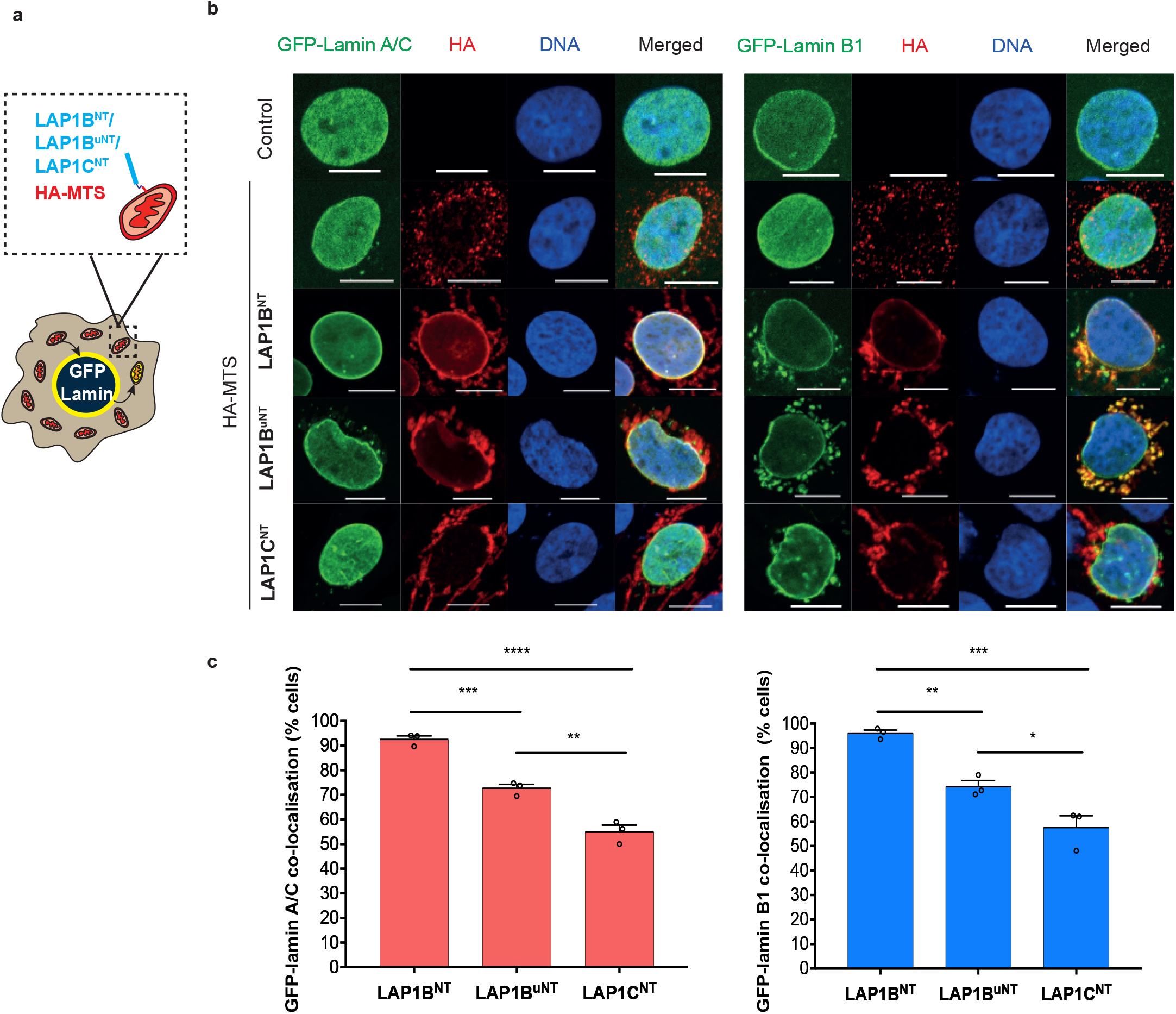
LAP1B and LAP1C are differentially tethered to nuclear lamins. **(a)** Schematic of mitochondrial retargeting assay. CAL51 cells were co-transfected with GFP-Lamin A/C or GFP-Lamin B1 and LAP1B^NT^/LAP1B^uNT^/LAP1C^NT^ tagged with HA-MTS. Where LAP1-Lamin interaction occurs, GFP-Lamin A/C or GFP-Lamin B1 would be dragged from the nuclear envelope to the mitochondria or LAP1B^NT^/LAP1B^uNT^/LAP1C^NT^ tagged with HA-MTS would be dragged from the mitochondria to the nuclear envelope. **(b)** Representative images of experimental results for retargeting assay in CAL51 cells expressing GFP-Lamin A/C or GFP-Lamin B1 and stained for HA (red) and DNA (blue). Scale bars, 5 μm. **(c)** Percentage of CAL51 cells showing co-localisation of GFP-Lamin A/C (left) or GFP-Lamin B1 (right) with LAP1B^NT^/LAP1B^uNT^/LAP1C^NT^. Left graph, n= 210, 211 and 190, respectively. Right graph, n= 211, 199 and 183, respectively. Experimental data have been pooled from three individual experiments. Graphs show the mean and error bars represent SEM. p values calculated by one-way ANOVA; *p < 0.05, **p < 0.01, ***p < 0.001, ****p < 0.0001.

Using cells stably expressing C-terminal mRuby3-tagged versions of LAP1C or LAP1B (M122A) (Supplementary Fig.7b), we observed that LAP1B-mRuby3, like GFP-Lamin B1, was largely excluded from NE blebs, whereas both LAP1C-mRuby3 and GFP-Lamin A/C were readily detectable in NE blebs (Supplementary Fig.7c, d). To determine if nuclear lamins were responsible for the differential incorporation LAP1 isoforms into NE blebs, we performed siRNA depletion of individual nuclear lamins (Supplementary Fig.7e). We found that endogenous LAP1 was present in NE blebs in the absence of Lamin B1 or Lamin B2, but not in the absence of Lamin A/C (Supplementary Fig.7f, g).

Lastly, using Fluorescence Recovery After Photobleaching (FRAP), we discovered that LAP1C-mRuby3 was more mobile than LAP1B-GFP both at the main NE and in NE blebs (Supplementary Fig.8 and Supplementary Movies 5a-c), in agreement with previous studies in other systems ^43^. We suggest that LAP1 isoforms are differentially mobile in the INM with LAP1B displaying stronger anchoring to chromatin and B-type lamins through its extended NT, whilst LAP1C can move more freely in the INM to populate NE blebs in a Lamin A/C dependent manner.

### LAP1C supports a nucleus permissive for blebbing and constrained migration

We next wondered if LAP1 isoforms made a differential contribution to constrained cell migration. In two-round transwell migration assays, we observed that stable expression of LAP1C-mRuby3 in WM983A cells increased both NE blebbing and migration efficiency (Fig.7a, b). Stable expression of both isoforms of LAP1 through LAP1-mRuby3 in WM983A cells similarly enhanced migratory capacity and NE blebbing (Fig.7a, b). In contrast, stable expression of LAP1B-mRuby3 in WM983A cells neither increased NE blebbing nor migration efficiency (Fig.7a, b). Furthermore, by sorting WM983A cells stably expressing LAP1C-mRuby3 according to mRuby expression levels we found that the effect of LAP1C promoting NE blebbing and constrained migration was concentration dependent (Fig.7c-g). We concluded that LAP1C is the dominant isoform allowing NE blebbing and conferring an ability for metastatic melanoma cells to migrate through multiple constraints.

**Figure 7.**
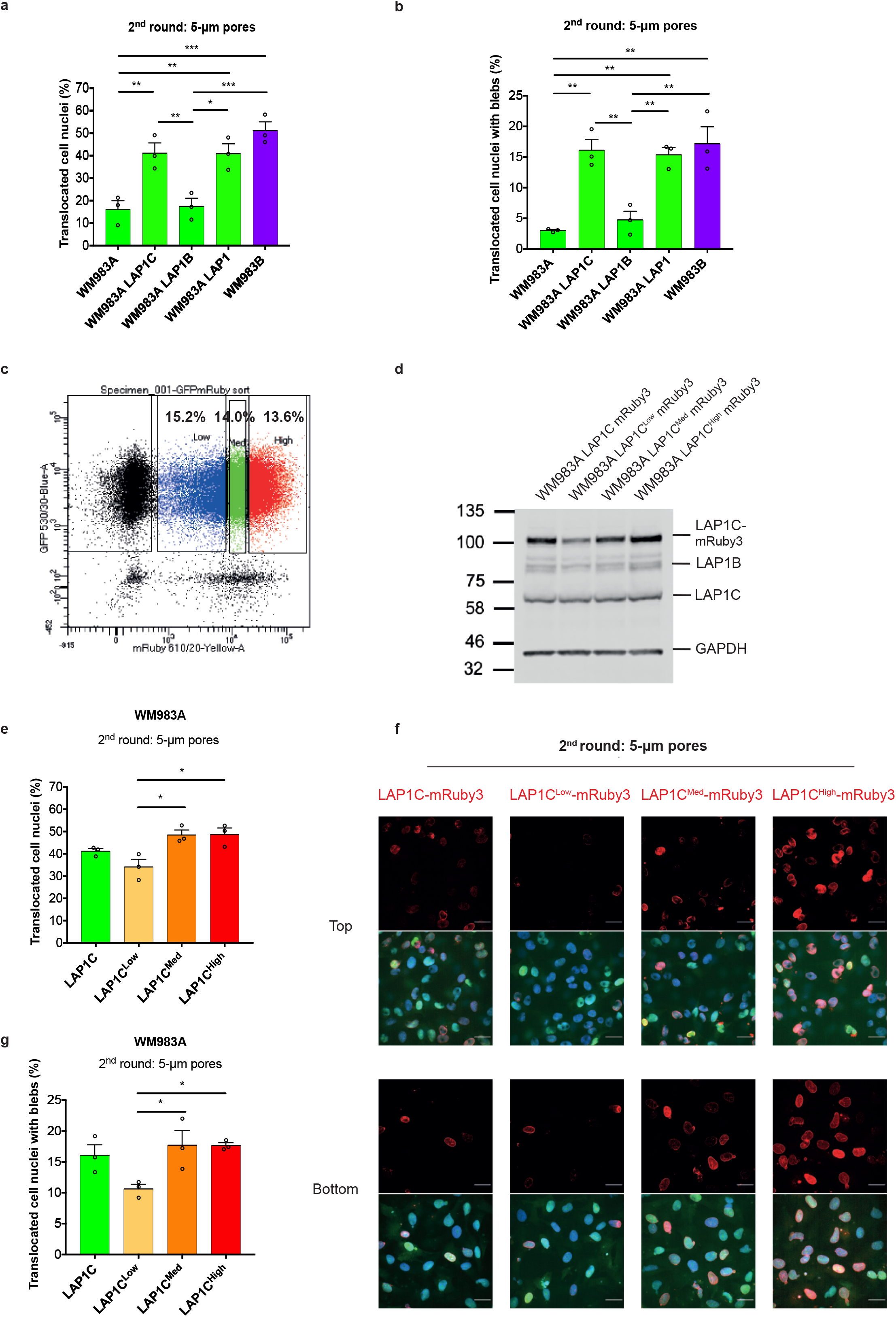
LAP1C supports a nucleus permissive for blebbing and constrained migration. **(a)** Percentage of primary melanoma WM983A cells, WM983A stably expressing LAP1C-mRuby3, LAP1B-mRuby3 (M122A) or LAP1-mRuby3 and metastatic melanoma WM983B cells that translocated their nuclei after a second round of transwell migration. **(b)** Percentage of WM983A cells, WM983A expressing LAP1C-mRuby3, LAP1B-mRuby3 (M122A) or LAP1-mRuby3 and WM983B cells that translocated their nuclei and displayed nuclear envelope blebs after a second round of transwell migration. n= 668, 694, 592, 722, 633, respectively. **(c)** FACS dot plot of WM983A cells stably expressing GFP-NLS and LAP1C-mRuby3 sorted according to levels of LAP1C-mRuby3 expression. **(d)** Representative immunoblot for endogenous and exogenous LAP1 expression levels in non-sorted WM983A cells and WM983A cells stably expressing GFP-NLS and LAP1C-mRuby3 sorted according to levels of LAP1C-mRuby3 expression. **(e)** Percentage of non-sorted WM983A cells and WM983A cells stably expressing GFP-NLS and LAP1C-mRuby3 sorted according to levels of LAP1C-mRuby3 expression that translocated their nuclei after a second round of transwell migration. **(f)** Representative pictures of non-sorted WM983A cells and WM983A cells stably expressing GFP-NLS (green) and LAP1C-mRuby3 (red) sorted according to levels of LAP1C-mRuby3 expression and stained for DNA (blue) after a second round of transwell migration. Scale bars, 30 μm. **(g)** Percentage of non-sorted WM983A cells and WM983A cells stably expressing GFP-NLS and LAP1C-mRuby3 sorted according to levels of LAP1C-mRuby3 expression that translocated their nuclei and displayed nuclear envelope blebs after a second round of transwell migration. n= 664, 601, 462, 531, respectively. Experimental data have been pooled from three individual experiments. Graphs show the mean and error bars represent SEM. p values calculated by one-way ANOVA; *p<0.05, **p< 0.01, ***p< 0.001.

### The differential tethers of LAP1 isoforms confer nuclear plasticity

Finally, we generated versions of LAP1B lacking either the dominant lamin-binding domain (LAP1B^Δ1-72^-mRuby3) or the CBR (LAP1B^ΔCBR^-mRuby3) (Fig.8a). Expression of LAP1B^Δ1-72^-mRuby3 or LAP1B^ΔCBR^-mRuby3 in WM983A cells enhanced both NE blebbing and migration in two-round transwell assays (Fig.8b-e), suggesting that an interplay between lamin and chromatin tethers restricts the nuclear shape rearrangements required for constrained migration. We validated the contribution of a truncated NT in constrained migration using siRNA resistant versions of wild-type and mutant LAP1B-mRuby3. Here, stable expression of LAP1B^Δ1-72^-mRuby3 or LAP1B^ΔCBR^-mRuby3, in a background of LAP1-depletion, allowed cells to generate NE blebs and migrate efficiently through a second constraint (Fig.8f, g).

**Figure 8.**
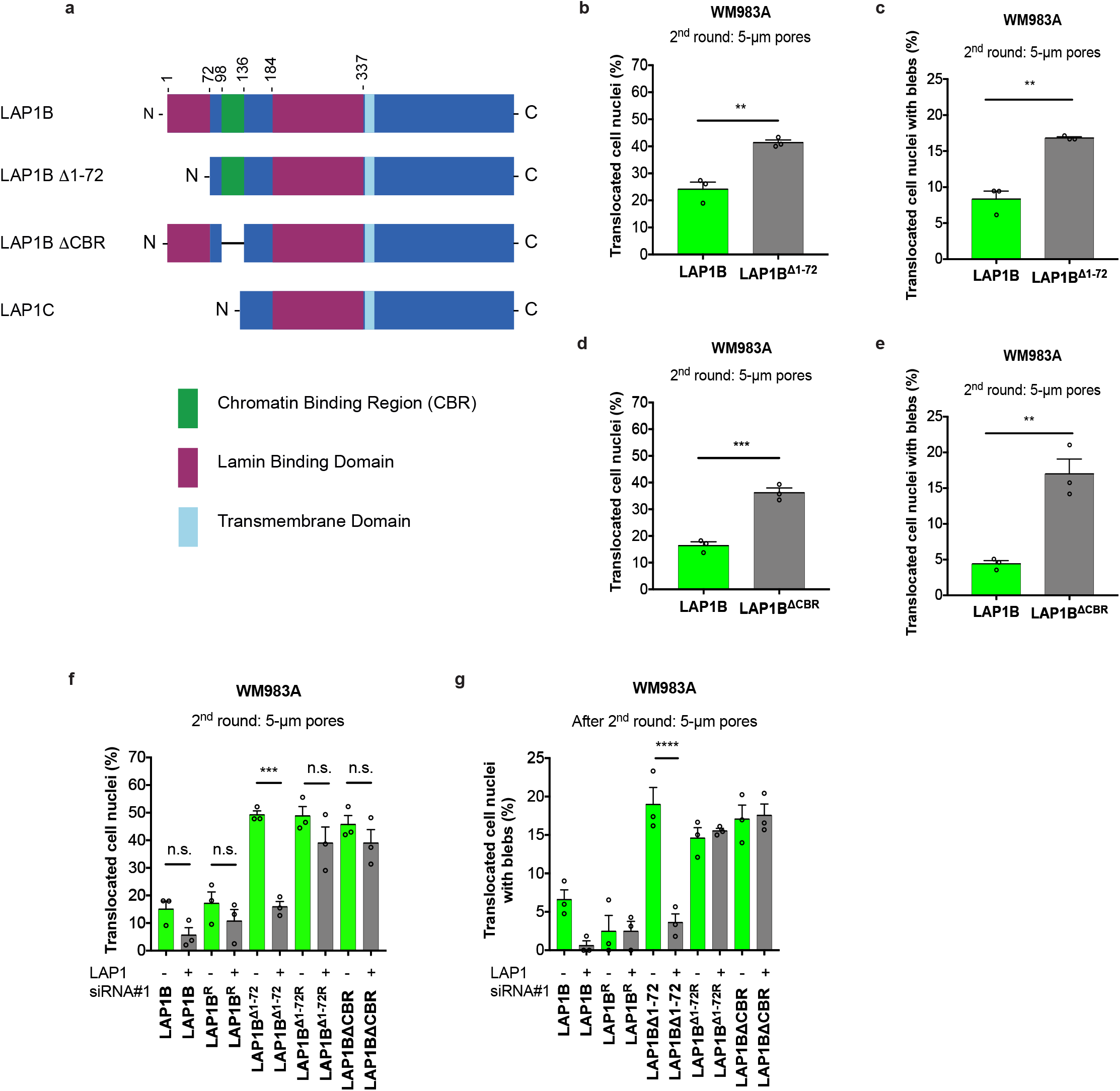
The differential tethers of LAP1 isoforms confer nuclear plasticity. **(a)** Schematic of LAP1B, LAP1B lacking lamin-binding region 1-72 (LAP1B^Δ1-72^), LAP1B lacking its chromatin-binding region (LAP1B^ΔCBR^) and LAP1C. Proteins have been drawn at aminoacid scale. **(b)** Percentage of primary melanoma WM983A cells stably expressing LAP1B-mRuby3 (M122A) or LAP1B^Δ1-72^-mRuby3 that translocated their nuclei after a second round of transwell migration. **(c**) Percentage of WM983A cells stably expressing LAP1B-mRuby3 (M122A) or LAP1B^Δ1-72^-mRuby3 that translocated their nuclei and displayed nuclear envelope blebs after a second round of transwell migration. n= 629 and 678, respectively. **(d)** Percentage of WM983A cells stably expressing LAP1B-mRuby3 (M122A) or LAP1B^ΔCBR^-mRuby3 that translocated their nuclei after a second round of transwell migration. **(e)** Percentage of WM983A cells stably expressing LAP1B-mRuby3 (M122A) or LAP1B^ΔCBR^-mRuby3 that translocated their nuclei and displayed nuclear envelope blebs after a second round of transwell migration. n= 518 and 545, respectively. **(f)** Percentage of WM983A stably expressing LAP1B-mRuby3 (M122A) or LAP1B^Δ1-72^-mRuby3, or resistant LAP1B-mRuby3 (LAP1B^R^-mRuby3), LAP1B^Δ1-72^-mRuby3 (LAP1B^Δ1-72R^-mRuby3) or LAP1B^ΔCBR^-mRuby3 that translocated their nuclei after a second round of transwell migration upon treatment with LAP1 siRNA#1. **(g)** Percentage of WM983A stably expressing LAP1B-mRuby3 (M122A), LAP1B^R^-mRuby3, LAP1B_Δ 1-72_-mRuby3, LAP1B^Δ1-72R^-mRuby3 or LAP1B^ΔCBR^-mRuby3 that translocated their nuclei and displayed nuclear envelope blebs after a second round of transwell migration upon treatment with LAP1 siRNA#1. n= 322, 269, 305, 357, 368, 363, 400, 369, 372 and 375, respectively. Experimental data have been pooled from three individual experiments. **a, b, d, e, f, g, h, i** Graphs show the mean and error bars represent SEM. p values calculated by one-way ANOVA and unpaired t-test; n.s.: not significant, **p< 0.01, ***p< 0.001, ****p < 0.0001.

Altogether, our data show how the distinct tethering of LAP1 isoforms to the lamina and chromatin enables metastatic melanoma cells to efficiently negotiate physical challenges. While LAP1B reinforces lamina and chromatin tethers, LAP1C supports nuclear deformability and licenses negotiation of physical constraints.

## DISCUSSION

Recent work recognises the importance of the cell nucleus during constrained migration ^45, 46^. Here, we describe that expression of *TOR1AIP1* and its encoded protein, the INM protein, LAP1, is upregulated in metastatic melanoma cells. Moreover, we observe that LAP1 is enriched at the IFs of melanoma tumours grown in mice. Using tissue sections from melanoma patients, we also found increased expression of LAP1 at the invasive front of primary melanomas and a further increase in expression in metastatic lesions. We find that LAP1 isoforms play distinct functions. For LAP1C, we find that its shorter NT lacking a lamin-binding domain and internal CBR supports NE blebbing and provides an advantage for migration through repeated constraints. Furthermore, we showed that the penetrance of this phenotype is directly related to LAP1C expression levels. On the other hand, we highlight that the longer NT of LAP1B, and specifically residues 1-72, enable a strong interaction with nuclear lamins, especially with lamin B1. LAP1B upregulation was reported to promote genetic instability ^43^. Another possible role for LAP1B, could be bringing chromatin closer to the NE to modulate gene expression. It is tempting to speculate that a stronger tethering to nuclear lamins could favour mechano-transduction ^47^.

NE blebs form and burst due to intra-nuclear hydrostatic pressure ^26, 27^. Whilst we report enhanced NE blebbing being associated with enhanced constrained migration, it is not clear whether blebs arise as a consequence of forcing a permissive nucleus through constraints, or whether the blebs themselves play an active role in negotiating constraints. Mechano-transduction at the cell nucleus relies on the connections between the nucleoskeleton and the cytoskeleton ^15, 18^. In melanoma cells, we describe that Rho-ROCK1/2-driven actomyosin contractility and LAP1C promote NE blebbing. We suggest that interactions with constraints *in vivo* might trigger a switch to high Rho-ROCK1/2-driven MLC2 activity in metastatic melanoma cells. We speculate that Rho-ROCK1/2-dependent transcriptional rewiring contributes to the nucleo-cytoskeletal rearrangements required for NE blebs to form and for the nucleus to translocate efficiently through subsequent constraints. Lamin A/C, LAP1C and potentially components of LINC complex and associated proteins ^48-50^ could participate in a bleb-based migratory programme favouring the establishment of nucleo-cytoskeletal interactions. This mode of migration relies upon hydrostatic pressure, to propel the nucleus through narrow spaces which could relive metastatic cells from mechanical stress and provide a survival advantage ^51^.

In the light of our results, we propose that by fine tuning the expression and localisation of LAP1 isoforms at the NE, metastatic melanoma cells acquire enhanced nuclear plasticity and gain an advantage for negotiating physical constraints. Thus, we propose that LAP1 could be a novel metastasis biomarker and that inhibiting LAP1 could be a route to prevent metastatic dissemination.

## ACKNOWLEDGMENTS

J.C.G. is a Wellcome Trust Senior Research Fellow (206346/Z/17/Z). V.S.-M. is a Cancer Research UK (CRUK) Senior fellow and V.S.M. lab was supported by (CRUK) C33043/A24478; Barts Charity; Fundación Alfonso Martin Escudero and Marie Sklodowska-Curie Action, grant agreement No 659022. Y.J.-G. received a Crick-KCL PhD studentship. We thank Dr. Eva Crosas-Molist and Dr. Jose L. Orgaz (Barts Cancer Institute, UK) for their supervision with cell biology experiments. We thank Professor Richard Marais (Cancer Research UK Manchester Institute) for the melanoma cells provided and Dr. Benilde Jiménez (Universidad Autónoma de Madrid and Instituto de Investigaciones Biomédicas CSIC-UAM, Spain) for the melanocytes. This work was supported in part by the Francis Crick Institute which receives its core funding from Cancer Research UK (FC001002), the UK Medical Research Council (FC001002), and the Wellcome Trust (FC001002). For the purpose of Open Access, the author has applied a CC BY public copyright licence to any Author Accepted Manuscript version arising from this submission.

## AUTHOR CONTRIBUTIONS

J.C.G. and V.S.-M. were principal investigators, designed the research, supervised experiments and wrote the paper; Y.J.-G. designed the research, performed the experiments, analysed the data, wrote the paper; O.M. assisted with the immunohistochemistry; I.R.-H. assisted with the transcriptomic analyses and qPCR; B.F. generated grew the tumours in mice; M.C.D. and L.M.C. performed and supervised the CLEM experiments; M.R. trained for and supervised the FRAP experiments; R.M.M. and X.M.-G provided the human tissue.

## MATERIALS AND METHODS

### Cell culture

The human melanoma cells WM1361 and WM1366 were from Professor Richard Marais (Cancer Research UK Manchester Institute); the human melanoma cells WM793B, WM983A, WM983B and WM88 were purchased from the Wistar Collection at Coriell Cell Repository; the human melanoma cells A375P and A375M2 were from Dr. Richard Hynes (HHMI, MIT); the human primary melanocytes M206 and M443 were a kind gift from Dr. Benilde Jiménez (Universidad Autónoma de Madrid and Instituto de Investigaciones Biomédicas CSIC-UAM, Spain) and were isolated from foreskins obtained with informed written consent from healthy donors and under approval of the Institutional Review Board of Hospital Infantil Universitario Niño Jesus (Madrid, Spain); HEK293T and CAL51 were from Dr. Jeremy Carlton (The Francis Crick Institute, UK). WM1361 and WM793B were cultured in Roswell Park Memorial Institute (RPMI, Gibco) medium supplemented with 10% of foetal bovine serum (FBS) and 1% (v/v) penicillin and streptomycin (PenStrep, Gibco); HEK293T, CAL51, WM1366, WM983A, WM983B, WM88, A375P and A375M2 were cultured in Dulbecco’s modified Eagle’s media (DMEM, Gibco) supplemented with 10% FBS and 1% (v/v) PenStrep; M206 and M443 were cultured in (MGM-4, Lonza) supplemented with 1 ml CaCl2, 2 ml BPE, 1 ml rhFGF-B, 1 ml rh-Insulin, 0.5 ml Hydrocortisone, 0.5 ml PMA, 0.5 ml GA-1000, 0.5% FBS and 1% (v/v) PenStrep. Cells were grown at 37°C and 5% or 10% CO_2_. Cells were kept in culture to a maximum of three or four passages.

### Plasmids

pTRIP-SFFV-EGFP-NLS and the coding sequences of mEmerald-Lamin A/C and mEmerald-Lamin B1 were from Addgene. pLVX-N-GFP was a kind gift from Prof. Michael Way (The Francis Crick Institute, UK) and was modified to express mEmerald-Lamin A/C or mEmerald-Lamin B1 by replacing GFP. The coding sequence for LAP1 was from Integrated DNA Technologies. LAP1C, point mutations in LAP1B (M122A, LAP1B^R^, LAP1B^Δ1-72R^) and deletion of lamin-binding residues 1-72 (Δ1-72) in LAP1B were generated using standard PCR. All PCR primers are in Supplementary Table 15. LAP1B carrying the deletion of the chromatin binding region (ΔCBR) was from Integrated DNA Technologies. LAP1, LAP1C and LAP1B mutants were cloned into pCMS28-EcoRI-NotI-XhoI-Linker-mRuby3. LAP1B (M122A) was also cloned into pNG72-EcoRI-NotI-XhoI-LAP-GFP. LAP1B^NT^, LAP1B^uNT^ or LAP1C^NT^ were cloned into pCR3.1 with HA-MTS.

### Transient transfection

CAL51 or WM983B cells were seeded at a density of 8×10^4^ cells in 4-well or 24-well plates and transfected with 0.5 μg of vector construct using optimem (Gibco), lipofectamine 3000 and P3000 (Invitrogen). Media was changed 6 hours post-transfection. At 48-hours post-transfection, CAL51 cells were fixed with 4% formaldehyde (FA) for 15 minutes at room temperature (RT).

### Generation of stable cell lines

HEK293T cells were seeded at a density of 4×10^5^ cells/ml in 6-well plates. For lentiviral production, cells were transfected with 1.5 μg of lentiviral vector, 0.5 μg of pVSVG and 2 μg of HIV-1 pCMVd8.91 using optimem, lipofectamine 3000 and P3000. For retroviral production, cells were transfected with 1.5 μg of retroviral vector, 0.5 μg of pHIT-VSVG and 2 μg of MLV-GagPol using optimem, lipofectamine 3000 and P3000. Media was changed 6 hours post-transfection. Viral supernatants were collected 48 hours post-transfection, spun down, and filtered (0.2 μm). Melanoma cells seeded at a density of 2×10^5^ in 6-well plates were infected with filtered supernatants. Antibiotic selection as required was started 48 hours post-infection. For assessing the concentration effect of LAP1C, cells were harvested in FACS buffer (PBS, 5mM EDTA, 2% FCS, 1mM HEPES) and sorted according to LAP1C-mRuby3 fluorescence intensity levels.

### siRNA transfection

Melanoma cells were seeded at a density of 4×10^4^ cells/ml in 24-well plates or 2.5×10^5^ in 6-well plates and transfected one hour after seeding. Cells were transfected with 20 mM siGenome Smart Pool or On-Targetplus LAP1 siRNA oligonucleotides using optimem and lipofectamine RNAimax (Invitrogen). All siRNA oligonucleotides were from Dharmacon and are listed in Supplementary Table 16. Non-targeting siRNA was used as a control. Two-round transwell migration and invasion assays were carried out 48-hours post-transfection.

### Transwell migration assays

Cells were starved in serum-free DMEM overnight and seeded at a density of 1.65×10^5^ cells/ml per insert in 24-well plates or 2.335×10^6^ cell/ml per insert in 6-well plates. DMEM 10% FBS was used as chemoattractant. Cells were allowed to migrate 16 hours in 24-well plates and 24 hours in 6-well plates. For multi-round transwell assays, cells were collected after one round from inserts in 6-well plates and seeded on inserts in 24-well plates. Cells were fixed with 4% FA for 15 minutes at RT. Transwell inserts were from Corning.

### Inhibitor treatments

ROCKi GSK269962A (Axon MedChem) was used at 1 μM and Staurosporin (Cell Guidance Systems) was used at 1 μM. In transwell assays, the inhibitor was added to the chemoattractant.

### Quantitative real-time PCR (qPCR)

Melanoma cells were seeded at a density of 1×10^5^ cells/ml in 12-well plates 24 hours prior to the experiment. RNA was extracted using Trizol (Life Technologies) following manufacturer’s instructions. RNA was treated with DNA-free™ DNA Removal Kit (Life Technologies) and RNA purity was determined with a ND-1000 Nanodrop (Thermo Fisher Scientific). qPCRs were performed using 100 ng RNA, QuantiTect primer assays and Brilliant III SYBR Green QRT-PCR Kit (Agilent Technologies) in a ViiA 7 Real-Time PCR System (Thermo Fisher Scientific). GAPDH was used as loading control. The following qPCR primers from Qiagen were used: RANBP2 (QT00035378), TPR (QT00046242), OSBPL8 (QT00067102), SUMO1 (QT00014280), NUP50 (QT00081669), ZMPSTE24 (QT00025627), TOR1AIP1 (QT00070147).

### Western blotting

Melanoma cells were seeded at a density of 6×10^5^ cells/ml in 12-well plates and lysed on the next day with LDS buffer 1X (Life Technologies). Lysates were denatured at 95°C for 5 minutes, sonicated and spun down. SDS-page electrophoresis was run at 150 V for 50 minutes. Transfer to PVDF membranes was run at 200 mA fixed for 90 minutes. Membranes were blocked for one hour in 5% BSA and incubated with primary antibody in 5% BSA overnight at 4°C in an orbital shaker. On the next day, membranes were incubated in secondary antibody in 5% BSA for 45 minutes at RT in an orbital shaker. Membranes were visualised using Odyssey Fc (LI-COR) and protein quantification was done with Odyssey Fc associated software. Primary antibodies were: LAP1 (1:1000; #21459-1-AP), Lamin A/C (1:1000, #10298-1-AP), Lamin B1 (1:1000, # 12987-1-AP) and Lamin B2 (1:1000, #10895-1-AP) from Proteintech; pThr18/Ser19-MLC2 (1:750, #3674), MLC2 (1:750, #3672) from Cell Signalling Technology; GAPDH (1:10,000, #MAB374) from Merck. Secondary antibodies were: IRDye 680RD goat anti-rabbit IgG (1:10,000, v925-68071) and IRDye 800RD goat anti-mouse IgG (1:10,000, #925-32210) from LI-COR.

### LAP1 solubilisation assay

The assays performed were an adaptation from ^39^. Protein solubilisation from melanoma cell lysates was carried out using the following buffers supplemented with 1 M DTT, 0.5 M NaF, 1 M glycerol phosphate, 100 mM VO_4_^-^, 0.1 M PMSF and ¼ EDTA tablet: 50 mM Tris-HCl pH 7.5; 50 mM Tris-HCl pH 7.5 + 1% Triton X-100; 50 mM Tris-HCl pH 7.5 + 1% Triton X-100 + 50 mM NaCl; 50 mM Tris-HCl pH 7.5 + 1% Triton X-100 + 500 mM NaCl. Protein lysates were incubated for 20 minutes on ice and then centrifuged at 14,000xg for 15 minutes. The supernatants were solubilised with LDS buffer 4X and the pellets with LDS buffer 1X, then they were denatured at 95°C for 5 minutes and western blotted.

### Cell culture on thick layers of collagen type I

Collagen I matrices were prepared using FibriCol (CellSystems) at 1.7 mg/ml. Collagen was left 4 hours to polimerise and melanoma cells were seeded on top at a density of 5×10^3^ in 96-well plates. Medium was changed the next day to DMEM 1% FBS. After 24 hours, cells were fixed with 20% FA for 15 minutes at RT. For nuclear staining, cells were fixed with 2% FA for 30 minutes at RT.

### 3D invasion assays

Collagen I was prepared using FibriCol at 2.3 mg/ml. Melanoma cells were suspended in serum-free DMEM to a final concentration of 1.5×10^4^ cells per 100 μl of collagen. Cells were centrifuged at 1800 rpm and 4°C for 8 minutes, resuspended in collagen and seeded on glass bottom 96-well plates (Ibidi). The plates were centrifuged at 900 rpm for 5 minutes to get all the cells at the bottom. Collagen was left 4 hours to polimerise and DMEM 10% FBS was added to the top. Cells were allowed to invade for 24 hours. Cells were fixed with 20% FA overnight at 4°C.

### Immunofluorescence staining

Cells in coverslips were fixed with 4% FA for 15 minutes at RT, permeabilised with 0.3% triton for 20 minutes, blocked with 4% bovine serum albumin (BSA) for 30 minutes and incubated sequentially with primary antibody in 4% BSA for two hours at RT and secondary antibody in 4% BSA for one hour at RT. DNA was stained with DAPI 1:1000 or Hoechst 1:5000 in PBS (Gibco). For DNA damage staining, cells were washed in PBS pre-fixation and incubated with CSK buffer (10 mM Pipes, pH 6.8, 100 mM NaCl, 300 mM sucrose, 3 mM MgCl_2_, 10mM B-glycerol phosphate, 50mM NaF, 1mM EDTA, 1mM EGTA, 5mM Na_2_VO_3_, 0.5% Triton) for three minutes and again for 1 minute on ice. Cells were fixed with 4% FA for 20 minutes on ice, blocked with 10% goat serum for one hour at RT and incubated sequentially with primary antibody in 1% goat serum overnight at 4°C and with secondary antibody in 1% goat serum for one hour at RT. For nuclear staining in collagen I, cells were permeabilised with 0.5% triton for 30 minutes, blocked with 4% BSA overnight at 4°C, incubated with primary antibody in 4% BSA overnight at 4°C and with secondary antibody in 4% BSA for two hours at RT. Primary antibodies were: Lamin A/C (1:200, #10298-1-AP) from Proteintech or (1:200; #mab3538) from Millipore; Lamin B1 (1:200, # 12987-1-AP), Lamin B2 (1:200, #10895-1-AP) and LAP1 (1:200; #21459-1-AP) from Proteintech; pSer19-MLC2 (1:200, #3671) from Cell Signalling); HA.11 (1:500, #901503) from BioLegend; Gamma-H2AX (1:600, #05-636) from Merck; 53BP1 (1:600, #NB100305) from Novus Biologicals. Secondary antibodies were Alexa Fluor 488 and Alexa Fluor 555 (1:1000) from Sigma or Invitrogen raised against the corresponding species. F-actin was stained using Alexa Fluor 546-phalloidin from Life Technologies and DNA with Hoechst 33342 from Invitrogen.

### Confocal fluorescence microscopy and image analysis

Images were acquired with a Dragonfly 200 high speed confocal microscope (Andor). Imaging was carried out with 20X or 60X oil objectives. Z-stacks of 1-5 μm step-size distance were acquired from fixed cells in 2D, transwells and collagen I. Live cell imaging was done at 37°C and 5% CO_2_ in a sealed chamber. Movies were taken at 2-minutes intervals for 1-15 hours. Image analysis was carried out using ImageJ. Fluorescence signal intensities were quantified from pixel intensity in single cells relative to the areas of interest.

### Fluorescence recovery after photobleaching

Cells were seeded at a density of 8×10^4^ in 4-well plates (Ibidi) 24 hours prior to the experiment. A LSM 880 Carl Zeiss microscope was used for imaging. Cells were kept at 37°C and 5% CO_2_ in a sealed chamber. Cells were imaged with a 63 × 1.4 NA objective. First, 5 pre-bleached values were acquired and then squares of 1.3 × 1.3 (1.69 μm^2^) at the main NE and at NE blebs were bleached with 50% 405 nm laser intensity. Fluorescence recovery was measured every second for 200 cycles. The FRAP data was curated subtracting background and normalised.

### Correlative light and electron microscopy

Melanoma cells were seeded at a density of 2×10^5^ in 35 mm gridded glass-bottom dishes (MatTek) 24 hours prior to the experiment. Live cells were imaged until nuclear envelope rupture was spotted, at which point cells were fixed adding 8 % (v/v) formaldehyde (Taab Laboratory Equipment Ltd, Aldermaston, UK) in 0.2 M phosphate buffer (PB) pH 7.4 to the cell culture medium (1:1) for 15 minutes. Cells were mapped using brightfield light microscopy to determine their position on the grid and tile scans were generated. The samples were then processed using a Pelco BioWave Pro+ microwave (Ted Pella Inc, Redding, USA) and following a protocol adapted from the National Centre for Microscopy and Imaging Research protocol ^52^. See Supplementary Table 17 for full BioWave program details. Each step was performed in the Biowave, except for the PB and water wash steps, which consisted of two washes on the bench followed by two washes in the Biowave without vacuum (at 250 W for 40 seconds). All the chemical incubations were performed in the Biowave for 14 minutes under vacuum in 2-minutes cycles alternating with/without 100W power. The SteadyTemp plate was set to 21°C unless otherwise stated. In brief, the samples were fixed again in 2.5% (v/v) gluteraldehyde (TAAB) / 4% (v/v) formaldehyde in 0.1M PB. The cells were then stained with 2% (v/v) osmium tetroxide (TAAB) / 1.5% (v/v) potassium ferricyanide (Sigma), incubated in 1% (w/v) thiocarbohydrazide (Sigma) with SteadyTemp plate set to 40°C, and further stained with 2% osmium tetroxide in ddH2O (w/v). The cells were then incubated in 1% aqueous uranyl acetate (Agar Scientific, Stansted, UK) with SteadyTemp plate set to 40°C, and then washed in dH_2_O with SteadyTemp set to 40°C. Samples were then stained with Walton’s lead aspartate with SteadyTemp set to 50°C, and dehydrated in a graded ethanol series (70%, 90%, and 100%, twice each), at 250 W for 40 seconds without vacuum. Exchange into Durcupan ACM® resin (Sigma) was performed in 50% resin in ethanol, followed by 4 pure Durcupan steps, at 250 W for 3 minutes, with vacuum cycling (on/off at 30-seconds intervals), before embedding at 60°C for 48 hours. Blocks were trimmed to a small trapezoid, excised from the resin block, and attached to a serial block-face scanning electron microscopy (SBF SEM) specimen holder using conductive epoxy resin. Prior to commencement of a SBF SEM imaging run, the sample were coated with a 2 nm layer of platinum to further enhance conductivity. SBF SEM data was collected using a 3View2XP (Gatan, Pleasanton, CA) attached to a Sigma VP SEM (Carl Zeiss Ltd, Cambridge, UK). Inverted backscattered electron images were acquired through the entire extent of the region of interest. For each of the 50-nm slices needed to image the cells in their whole volume, a low-resolution overview image (horizontal frame width 103 µm; pixel size of 40 nm; using a 2 µseconds dwell time) and a high-resolution image of the cell of interest (horizontal frame width 32 and 39 µm respectively; pixel size of 8 nm; using a 2 µseconds dwell time) were acquired. The SEM was operated in high vacuum with focal charge compensation on (70%). The 30 µm aperture was used, at an accelerating voltage of 1.8 kV. Only minor adjustments in image alignment were needed and were done using the TrakEM2 plug-in of the FIJI framework ^53^.

### Immunohistochemistry

Whole sections from subcutaneous tumours (A375P/A375M2, WM983A/WM983B) and two tissue microarrays including a total of 48 primary melanoma tumours and 43 metastases, were used. Patient biopsies were obtained with approvals from the Ethics and Scientific Committee and specific informed patient consent ^32^. The IFs were delimited as the tumour areas showing at least 50% contact with the matrix, as previously described ^5-7, 32^. All samples were formalin-fixed paraffin-embedded tissue. Samples were sectioned (3-4 μm thick) and dried for one hour at 65°C. Next, samples were deparaffined and rehydrated and endogenous peroxidase activity was blocked with 3% H_2_O_2_ in ethanol absolute for 10 minutes. Heat-induced epitope retrieval was carried out using 1:100 pH 6 Citrate Buffer H-3300 for 10 minutes at 100°C in a Biocare Decloaking Chamber (DC2012). Incubation with primary antibody in Zytomed antibody diluent was carried out for 40 minutes. Incubation with secondary antibody polymer conjugated (ImmPRESS Polymer Reagent) was carried out for 45 minutes. Incubation with Vector VIP HRP substrate chromogen was done for up to 10 minutes. All reagents used for detection were from VECTASTAIN ABC-HRP Kit (PK-4000). All samples were counterstained with haematoxylin. Lastly, samples were dehydrated, and slides were mounted. Reagents were used at RT in humidified slide chambers. Primary antibodies were: LAP1 (1:100; #21459-1-AP) from Proteintech; melanA (1:100, #ab51061) from Abcam. Whole section images were obtained from each sample using a NanoZoomer S210 slide scanner (Hamamatsu, Japan). Image analysis was done using QuPath software ^54^. Positive cell detection was carried out and threshold to the intensity scores (0,1, 2, 3) was applied. Then, QuPath software was trained to differentiate tumour cells from stroma, staining was graded semiquantitatively and H-scores were calculated as previously described ^32^. Co-localisation analysis was conducted for LAP1 and melanA staining as previously described ^32^.

### Gene enrichment analysis

Normalised gene expression microarray data was obtained from GSE23764 ^6^. A375M2 were compared to A375P and to A375M2 treated 24 hours with contractility inhibitors (ROCK inhibitors H1152 or Y27632 or myosin inhibitor blebbistatin). A catalogue of nuclear gene sets was downloaded, and analyses were carried out using GSEA software (http://www.broadinstitute.org/gsea/index.jsp). Permutations were set to 1000, permutation type to gene-set and t-test was established for ranking. GO gene sets were classified according to GO ontologies. Upregulated gene sets were filtered based on FDR<5% and p-value<0.05.

### Gene expression analyses

Gene expression data from cells was obtained from publicly available datasets and normalised as previously described ^35, 55^. Data from 28 melanocytes was obtained from four melanocyte datasets from ^35^ (GSE4570, GSE4840), ^56^ and ^57^, and data from melanoma cell lines (Philadelphia cohort GSE4841 (29 samples) and Mannheim cohort GSE4843 (37 samples)) where obtained from ^35^. Heatmaps for gene expression in cells were generated using MeV_4_9_0 software (http://mev.tm4.org/). Gene expression data from human samples was derived from three publicly available datasets from ^37^ (Riker GSE755333 (14 primary and 40 metastatic melanomas)), ^36^ (Kabbarah GSE4651732 (31 primary and 73 metastatic melanomas)) and ^38^(Xu GSE840134 (31 primary and 52 metastatic melanomas)).

### Statistical analyses

Unpaired t test, one-way ANOVA and two-way ANOVA followed by Tukey’s or Sidak’s multiple comparisons tests as appropriate were performed using GraphPad Prism (GraphPad Software, Inc). All results were obtained from at least three independent experiments unless otherwise stated. Data is plotted as minimum to maximum boxplots or graphs with the mean ± standard error of the mean (SEM). The threshold for statistical significance was set to a p-value of less than 0.05.

