## Supplemental Text for "LAP1 supports nuclear plasticity during constrained migration"

**SUPPLEMENTARY FIGURE LEGENDS**

**Supplementary Figure 1. Multi-round transwell assays of melanoma cells**

**(a)** Percentage of primary melanoma WM983A and metastatic melanoma WM983B cells that did not translocate their nuclei after a first round of migration in transwells of various pore sizes. **(b)** Percentage of WM983A and WM983B cells that did not translocate their nuclei and displayed nuclear envelope blebs after a first round of migration in transwells of various pore sizes. **(c)** Percentage of WM983A and WM983B cells that translocated their nuclei and displayed nuclear envelope blebs classified according to bleb number and bleb shape after a first round of migration in transwells of various pore sizes. n= 1758 and 1728, respectively. **(d)** Percentage of WM983A and WM983B cells that did not translocate their nuclei after a second round of transwell migration. **(e)** Percentage of WM983A and WM983B cells that did not translocate their nuclei and displayed nuclear envelope blebs after a second round of transwell migration. **(f)** Percentage of WM983A and WM983B cells that translocated their nuclei and displayed nuclear envelope blebs classified according to bleb number and bleb shape after a second round of transwell migration. n= 192 and 313, respectively. **(g)** Percentage of WM983A and WM983B cells that translocated their nuclei after a third round of transwell migration. **(h)** Percentage of WM983A and WM983B cells that did not translocate their nuclei after a third round of transwell migration. **(i)** Percentage of WM983A and WM983B cells that translocated their nuclei and displayed nuclear envelope blebs after a third round of transwell migration. **(j)** Relative percentage of WM983A and WM983B cells that did not translocate their nuclei and displayed nuclear envelope blebs after a third round of transwell migration. n= 145 and 170, respectively. N=1. Experimental data have been pooled from three individual experiments unless otherwise stated. Graphs show the mean and error bars represent SEM. p values calculated by one-way ANOVA, two-way ANOVA, and unpaired t test; *p < 0.05, **p< 0.01, ***p< 0.001, ****p < 0.0001.

**Supplementary Figure 2. Cell viability and actomyosin contractility in multi-round transwell assays**

**(a)** Percentage of alive primary melanoma WM983A cells and metastatic melanoma WM983B cells after a first round of migration in transwells. **(b)** Percentage of alive WM983A and WM983B cells after a second round of migration in transwells upon staurosporin treatment. n= 180 and 213, respectively. N=1. **(c)** Representative pictures of WM983A and WM983B cells expressing GFP-NLS (green) and stained for caspase-3 (red) and DNA (blue) after a second round of migration in transwells upon staurosporin treatment. Scale bars, 30 μm. **(d)** Representative immunoblot for WM983A and WM983B cells after treatment with ROCK inhibitor (ROCKi) GSK269962A. **(e)** Representative pictures of WM983A and WM983B cells treated with ROCKi and stained for pMLC2 (red), lamin A/C (green) and DNA (blue) after a first round of transwell migration. Scale bars, 30 μm. Experimental data have been pooled from three individual experiments unless otherwise stated. Graphs show means and error bars represent SEM. p values calculated by unpaired t test. n.s.: not significant.

**Supplementary Figure 3. Nuclear shape changes in melanoma progression**

**(a)** Percentage of nuclei with nuclear envelope blebs in M206 melanocytes. n= 120. N=1. **(b)** Representative picture of M206 melanocytes stained for lamin A/C (red) and DNA (blue). Scale bars, 30 μm. The magnification shows a M206 nucleus with a nuclear envelope bleb. Scale bar, 10 μm. **(c)** Cell roundness index, **(d)** percentage of elongated and rounded cells and **(e)** pMLC2 levels in primary melanoma WM983A cells and metastatic melanoma WM983B cells on collagen I. n= 375 and 425, respectively. **(f)** Cell roundness index, **(g)** percentage of elongated and rounded cells and **(h)** pMLC2 levels in less metastatic melanoma A375P cells and highly metastatic melanoma A375M2 cells on collagen I. n= 385 and 381, respectively. **(i)** Representative images of WM983A, WM983B, A375P and A375M2 cells stained for pMLC2 (green), actin (red) and DNA (blue) on collagen I. Scale bars, 30 μm. The magnifications show representative cells. Scale bars, 10 μm. **(j)** Percentage of nuclei with nuclear envelope blebs, **(k)** nuclear circularity and **(l)** karyoplasmic ratio in WM983A and WM983B cells on collagen I. n= 337 and 373, respectively. **(m)** Percentage of nuclei with nuclear envelope blebs, **(n)** nuclear circularity and **(o)** karyoplasmic ratio in A375P and A375M2 cells on collagen I. n= 335 and 331, respectively. **(p)** Representative images of WM983A, WM983B, A375P and A375M2 cells stained for lamin A/C (green), actin (red) and DNA (blue) on collagen I. Scale bars 30 μm. The magnifications show representative cells. Scale bars, 10 μm. Experimental data have been pooled from three individual experiments. **a, d, g, j, m** Graphs show the mean and error bars represent SEM. **c, e, f, h, k, l, n, o** Horizontal lines show the median and whiskers show minimum and maximum range of values. p values calculated by two-way ANOVA and unpaired t test; **p < 0.01, ***p < 0.001, ****p < 0.0001.

**Supplementary Figure 4. Instability of nuclear envelope blebs in melanoma cells**

**(a)** Percentage of nuclear envelope blebs positive for DNA damage markers (53BP1 and gamma-H2AX) in primary melanoma WM983A cells and metastatic melanoma WM983B cells. **(b)** Representative pictures of a WM983B nucleus with a nuclear envelope bleb positive for DNA damage stained for Gamma-H2AX (green), 53BP1 (red) and DNA (blue). Scale bars, 10 μm. **(c)** Relative Gamma-H2AX and 53BP1 fluorescence intensity levels and average Gamma-H2AX and 53BP1 foci per nucleus in WM983A and WM983B cells with or without nuclear envelope blebs. n= 246 and 283, respectively. N=2. **(d)** Representative image sequence from repetitive, transient nuclear envelope rupture events in WM983A nucleus over 5 hours. **(e)** Representative image sequence from repetitive, transient nuclear envelope rupture events in a WM983B nucleus over 5 hours. Images showing the first frame of a nuclear envelope rupture in (d) or (e) are surrounded by a red square. Scale bars, 10 μm. Experimental data have been pooled from three individual experiments unless otherwise stated. Graphs show the mean and error bars represent SEM.

**Supplementary Figure 5. LAP1 expression in tissue**

**(a)** Karyoplasmic ratio of tumour cells according to LAP1 intensity score in tumour bodies (TB) and invasive fronts (IF) of WM983A and WM983B melanoma tumours grown in SCID mice. n= 8 and 4, respectively. **(b)** Karyoplasmic ratio of tumour cells according to LAP1 intensity score in TBs and IFs of A375P and A375M2 melanoma tumours grown in SCID mice. n= 8 and 8, respectively. **(c)** Representative pictures of normal skin stained for LAP1 (pseudocoloured in magenta), melanA (pseudocoloured in green) and DNA (pseudocoloured in blue). Scale bars, 50 μm and 30 μm, magnifications. **(d)** Levels of LAP1 measured in melanocytes and keratinocytes from pseudocoloured images. n= 20 and 20, respectively. **(e)** Karyoplasmic ratio of tumour cells according to LAP1 intensity score in TBs and IFs of primary tumours and metastases of human melanoma patient cohort A. n= 19 and 14, respectively. **(f)** Karyoplasmic ratio of tumour cells according to LAP1 intensity score in TBs and IFs of primary tumours and metastases of human melanoma patient cohort B. n= 29 and 29, respectively. **d** Horizontal lines show the median and whiskers show minimum and maximum range of values. p value calculated by two-way ANOVA and unpaired t test; ****p < 0.0001.

**Supplementary Figure 6. Impact of LAP1 siRNA on LAP1 expression, melanoma cell viability and migration**

**(a)** Quantification of LAP1B and LAP1C expression in metastatic melanoma WM983B cells after 48 hours of LAP1 depletion with a siGENOME SMARTpool. **(b)** Percentage of alive WM983B cells after a first round of transwell migration upon LAP1 depletion with a siGENOME SMARTpool. **(c)** Representative immunoblot for LAP1 expression levels in WM983B cells after 48 hours of LAP1 depletion with ON-TARGETplus individual siRNAs. **(d)** Quantification of LAP1B and LAP1C expression in WM983B cells after 48 hours of LAP1 depletion with ON-TARGETplus individual siRNAs. **(e)** Percentage of WM983B cells that translocated their nuclei after a second round of transwell migration upon LAP1 depletion with ON-TARGETplus individual siRNAs. **(f)** Percentage of WM983B cells that translocated their nuclei and displayed nuclear envelope blebs after a second round of transwell migration upon LAP1 depletion with ON-TARGETplus individual siRNAs. n= 772, 632 and 590, respectively. Experimental data have been pooled from three individual experiments. Graphs show the mean and error bars represent SEM. p values calculated by one-way ANOVA and unpaired t test; n.s.: not significant, *p < 0.05, **p < 0.01, ***p < 0.001, ****p < 0.0001.

**Supplementary Figure 7. Lamin-binding properties of LAP1 isoforms**

**(a)** LAP1 solubilisation assay in metastatic melanoma WM983B cells. S: supernatant; P: pellet. **(b)** Representative immunoblot for endogenous and exogenous LAP1 expression levels in primary melanoma WM983A cells, WM983A stably expressing LAP1C-mRuby3, LAP1B-mRuby3 (M122A) or LAP1-mRuby3 (allowing expression of both LAP1 isoforms) and metastatic melanoma WM983B cells. **(c)** Representative pictures of WM983B transfected with GFP-lamin A/C or GFP-lamin B1 and stably expressing LAP1B-mRuby3 (M122A) or LAP1C-mRuby3 displaying nuclear envelope blebs. Scale bars, 10 μm. **(d)** Levels of LAP1B-mRuby3 (M122A), LAP1C-mRuby3, GFP-LAMIN A/C and GFP-LAMIN B1 measured in WM983B cells. n= 43, 63, 30 and 42, respectively. **(e)** Representative immunoblot for lamin A/C, LAP1, lamin B1 and lamin B2 expression levels in WM983B cells after 72 hours of siRNA depletion of lamin A/C, lamin B1 or lamin B2 with siGENOME SMARTpools. **(f)** Representative pictures of WM983B stably expressing GFP-NLS (green) stained for LAP1 (red) and DNA (blue). Scale bars, 10 μm. **(g)** Percentage of nuclear envelope blebs containing LAP1 upon siRNA depletion of lamin A/C, lamin B1 or lamin B2 with siGENOME SMARTpools. Experimental data have been pooled from three individual experiments. Graphs show the mean and error bars represent SEM. p values calculated by unpaired t tests; n.s.: not significant, *p < 0.05, ****p < 0.0001.

**Supplementary Figure 8. Motility of LAP1 isoforms at the NE and into NE blebs**

**(a)** Representative image sequence of FRAP experiment in metastatic melanoma WM983B cells stably co-expressing LAP1B-GFP (M122A) (green) and LAP1C-mRuby3 (red). Scale bars, 5 μm. The white arrowheads indicate where the cell was bleached at the main nuclear envelope and at the nuclear envelope bleb. **(b)** FRAP analysis of the relative motility of LAP1 isoforms at the main nuclear envelope in WM983B cells stably co-expressing LAP1B-GFP (M122A) and LAP1C-mRuby3. **(c)** FRAP analysis of the relative motility of LAP1 isoforms at nuclear envelope blebs in WM983B cells stably co-expressing LAP1B-GFP (M122A) and LAP1C-mRuby3. n= 26. Experimental data have been pooled from three individual experiments. Graphs show the mean and error bars represent SEM.

**SUPPLEMENTARY MOVIE LEGENDS**

**Supplementary Movie 1A. Intact nuclear envelope bleb by SBF SEM.**

Representative SBF SEM reconstruction of the nucleus of a metastatic melanoma WM983B cell with an intact nuclear envelope bleb. Scale bar, 5 μm.

**Supplementary Movie 1B. Intact nuclear envelope bleb by CLEM.**

Representative CLEM movie of the nucleus of a metastatic melanoma WM983B cell stably expressing GFP-NLS (green) with an intact nuclear envelope bleb. Time interval: 6 min. Scale bar, 5 μm.

**Supplementary Movie 2A. Ruptured nuclear envelope bleb by SBF SEM.**

Representative SBF SEM reconstruction of the nucleus of a metastatic melanoma WM983B cell with a ruptured nuclear envelope bleb. Scale bar, 5 μm.

**Supplementary Movie 2B. Ruptured nuclear envelope bleb by CLEM.**

Representative CLEM movie of the nucleus of a metastatic melanoma WM983B cell stably expressing GFP-NLS (green) with a ruptured nuclear envelope bleb. Time interval: 6 min. Scale bar, 5 μm.

**Supplementary Movie 3. Nuclear envelope rupture in primary melanoma cell.**

Representative movie of nuclear envelope rupture in a WM983A cell stably expressing GFP-NLS (green). Time interval: 5 hours. Scale bar, 10 μm.

**Supplementary Movie 4. Nuclear envelope rupture in metastatic melanoma cell.**

Representative movie of nuclear envelope rupture in a WM983B cell stably expressing GFP-NLS (green). Time interval: 5 hours. Scale bar, 10 μm.

**Supplementary Movie 5A. FRAP of LAP1 isoforms in metastatic melanoma cell.**

Representative movie of FRAP in a WM983B nucleus stably co-expressing LAP1B-GFP (M122A) (green) and LAP1C-mRuby3 (red) and with a nuclear envelope bleb. FRAP was measured at the main nuclear envelope and at the bleb. Time interval: 200 seconds. Scale bar, 5 μm.

**Supplementary Movie 5B. FRAP of LAP1B in metastatic melanoma cell.**

Representative movie of FRAP in a WM983B nucleus stably co-expressing LAP1B-GFP (M122A) (green) and LAP1C-mRuby3 (red) and with a nuclear envelope bleb. FRAP was measured at the main nuclear envelope and at the bleb. Time interval: 200 seconds. Scale bar, 5 μm.

**Supplementary Movie 5C. FRAP of LAP1C in metastatic melanoma cell.**

Representative movie of FRAP in a WM983B nucleus stably co-expressing LAP1B-GFP (M122A) (green) and LAP1C-mRuby3 (red) and with a nuclear envelope bleb. FRAP was measured at the main nuclear envelope and at the bleb. Time interval: 200 seconds. Scale bar, 5 μm.
