## Supplemental Table 1-17 for "LAP1 supports nuclear plasticity during constrained migration"

### Supplementary Table 1. Signalling regulation gene sets enriched in A375M2.

| Signalling regulation | | | | | | | |
|---|---|---|---|---|---|---|---|
| **GS follow link to MSigDB** | **SIZE** | **ES** | **NES** | **NOM p-val** | **FDR q-val** | **FWER p-val** | **RANK AT MAX** |
| GO_MACROMOLECULAR_COMPLEX_BINDING | 1236 | 0.33 | 1.59 | 0.000 | 0.005 | 0.102 | 3831 |
| GO_REGULATION_OF_PROTEIN_COMPLEX_DISASSEMBLY | 179 | 0.36 | 1.56 | 0.001 | 0.006 | 0.128 | 3222 |
| GO_NEGATIVE_REGULATION_OF_PROTEIN_COMPLEX_DISASSEMBLY | 140 | 0.36 | 1.52 | 0.005 | 0.008 | 0.191 | 2941 |
| GO_NEGATIVE_REGULATION_OF_PHOSPHORYLATION | 381 | 0.35 | 1.59 | 0.000 | 0.009 | 0.069 | 3711 |
| GO_CELLULAR_MACROMOLECULE_LOCALIZATION | 1079 | 0.31 | 1.48 | 0.000 | 0.011 | 0.282 | 4335 |
| GO_PROTEIN_COMPLEX_BINDING | 839 | 0.31 | 1.47 | 0.000 | 0.012 | 0.291 | 3831 |
| GO_PROTEIN_COMPLEX_SUBUNIT_ORGANIZATION | 1324 | 0.28 | 1.35 | 0.000 | 0.029 | 0.685 | 4644 |
| GO_NEGATIVE_REGULATION_OF_PHOSPHORUS_METABOLIC_PROCESS | 478 | 0.32 | 1.46 | 0.000 | 0.017 | 0.209 | 3711 |
| GO_NEGATIVE_REGULATION_OF_CELL_COMMUNICATION | 1066 | 0.29 | 1.38 | 0.000 | 0.028 | 0.396 | 3267 |
| GO_REGULATION_OF_PROTEIN_KINASE_B_SIGNALING | 111 | 0.35 | 1.38 | 0.041 | 0.027 | 0.396 | 3082 |
| GO_NEGATIVE_REGULATION_OF_PROTEIN_MODIFICATION_PROCESS | 539 | 0.30 | 1.38 | 0.003 | 0.026 | 0.403 | 3714 |
| GO_NEGATIVE_REGULATION_OF_MAPK_CASCADE | 133 | 0.33 | 1.36 | 0.038 | 0.028 | 0.459 | 4921 |
| GO_NEGATIVE_REGULATION_OF_INTRACELLULAR_SIGNAL_TRANSDUCTION | 394 | 0.29 | 1.36 | 0.009 | 0.028 | 0.476 | 3249 |
| GO_NEGATIVE_REGULATION_OF_RESPONSE_TO_STIMULUS | 1215 | 0.28 | 1.35 | 0.000 | 0.028 | 0.488 | 3275 |
| GO_REGULATION_OF_PROTEIN_MODIFICATION_PROCESS | 1502 | 0.28 | 1.35 | 0.000 | 0.029 | 0.509 | 3621 |
| GO_NEGATIVE_REGULATION_OF_PROTEIN_METABOLIC_PROCESS | 946 | 0.27 | 1.31 | 0.004 | 0.039 | 0.625 | 3726 |
| GO_REGULATION_OF_INTRACELLULAR_SIGNAL_TRANSDUCTION | 1476 | 0.27 | 1.31 | 0.000 | 0.039 | 0.638 | 3809 |
| GO_ENZYME_BINDING | 1541 | 0.32 | 1.56 | 0.000 | 0.011 | 0.057 | 4342 |
| GO_NUCLEOSIDE_TRIPHOSPHATASE_REGULATOR_ACTIVITY | 258 | 0.42 | 1.84 | 0.000 | 0.013 | 0.011 | 3581 |
| GO_ATPASE_REGULATOR_ACTIVITY | 28 | 0.58 | 1.78 | 0.005 | 0.010 | 0.017 | 3138 |
| GO_ENZYME_ACTIVATOR_ACTIVITY | 384 | 0.35 | 1.58 | 0.000 | 0.015 | 0.087 | 4329 |
| GO_POSITIVE_REGULATION_OF_HYDROLASE_ACTIVITY | 768 | 0.32 | 1.53 | 0.000 | 0.015 | 0.131 | 4341 |
| GO_ATPASE_ACTIVATOR_ACTIVITY | 15 | 0.57 | 1.51 | 0.055 | 0.016 | 0.157 | 3059 |
| GO_POSITIVE_REGULATION_OF_MOLECULAR_FUNCTION | 1564 | 0.30 | 1.45 | 0.000 | 0.020 | 0.222 | 3607 |
| GO_POSITIVE_REGULATION_OF_CATALYTIC_ACTIVITY | 1315 | 0.30 | 1.44 | 0.000 | 0.021 | 0.248 | 4407 |
| GO_POSITIVE_REGULATION_OF_ATPASE_ACTIVITY | 37 | 0.43 | 1.41 | 0.062 | 0.021 | 0.291 | 3059 |
| GO_ENZYME_REGULATOR_ACTIVITY | 804 | 0.28 | 1.32 | 0.002 | 0.042 | 0.518 | 3606 |
| GO_MOLECULAR_FUNCTION_REGULATOR | 1161 | 0.27 | 1.31 | 0.000 | 0.045 | 0.571 | 3606 |
| GO_POLY_A_RNA_BINDING | 938 | 0.37 | 1.78 | 0.000 | 0.005 | 0.011 | 5503 |
| GO_RNA_BINDING | 1242 | 0.36 | 1.74 | 0.000 | 0.006 | 0.019 | 4777 |
| GO_PROTEIN_DOMAIN_SPECIFIC_BINDING | 563 | 0.35 | 1.63 | 0.000 | 0.008 | 0.040 | 4151 |
| GO_OXIDOREDUCTASE_ACTIVITY_ACTING_ON_THE_CH_CH_GROUP_OF_DONORS_NAD_OR_NADP_AS_ACCEPTOR | 21 | 0.55 | 1.60 | 0.031 | 0.010 | 0.060 | 1886 |

#### Supplementary Table 2. Nuclear membrane gene sets enriched in A375M2.

| Nuclear membrane | | | | | | | |
|---|---|---|---|---|---|---|---|
| GS follow link to MSigDB | SIZE | ES | NES | NOM p-val | FDR q-val | FWER p-val | RANK AT MAX |
| GO_NUCLEAR_MEMBRANE | 233 | 0.38 | 1.68 | 0.000 | 0.003 | 0.039 | 3805 |
| GO_NUCLEAR_ENVELOPE | 349 | 0.34 | 1.54 | 0.000 | 0.007 | 0.151 | 4522 |
| GO_NUCLEAR_ INNER_MEMBRANE | 39 | 0.46 | 1.50 | 0.039 | 0.010 | 0.243 | 3902 |
| GO_ENDOPLASMIC_ RETICULUM | 1336 | 0.26 | 1.28 | 0.001 | 0.046 | 0.223 | 3811 |
| GO_ENDOPLASMIC_ RETICULUM_PART | 951 | 0.27 | 1.27 | 0.006 | 0.045 | 0.250 | 3811 |
| GO_NUCLEAR_PERIPHERY | 111 | 0.51 | 2.05 | 0.000 | 0.000 | 0.000 | 4367 |

#### Supplementary Table 3. Cell division gene sets enriched in A375M2.

| Cell division | | | | | | | |
|---|---|---|---|---|---|---|---|
| GS follow link to MSigDB | SIZE | ES | NES | NOM p-val | FDR q-val | FWER p-val | RANK AT MAX |
| GO_MITOTIC_CELL_CYCLE | 643 | 0.44 | 2.07 | 0.000 | 0.000 | 0.000 | 4141 |
| GO_CHROMOSOMAL_ REGION | 266 | 0.46 | 2.07 | 0.000 | 0.000 | 0.000 | 4648 |
| GO_CELL_CYCLE | 1097 | 0.42 | 2.05 | 0.000 | 0.000 | 0.000 | 3676 |
| GO_CELL_CYCLE_PROCESS | 900 | 0.42 | 2.04 | 0.000 | 0.000 | 0.000 | 4159 |
| GO_CONDENSED_ CHROMOSOME | 160 | 0.48 | 2.02 | 0.000 | 0.000 | 0.001 | 3830 |
| GO_CHROMOSOME | 736 | 0.42 | 2.00 | 0.000 | 0.000 | 0.001 | 3830 |
| GO_MICROTUBULE_BASED_ PROCESS | 420 | 0.40 | 1.82 | 0.000 | 0.001 | 0.006 | 3916 |
| GO_REGULATION_OF_ MICROTUBULE_BASED_ PROCESS | 194 | 0.42 | 1.80 | 0.000 | 0.001 | 0.006 | 3271 |
| GO_TUBULIN_BINDING | 229 | 0.40 | 1.77 | 0.000 | 0.001 | 0.011 | 3845 |
| GO_MITOTIC_SPINDLE_ ORGANIZATION | 57 | 0.50 | 1.77 | 0.002 | 0.001 | 0.011 | 3881 |
| GO_MICROTUBULE_BINDING | 171 | 0.40 | 1.69 | 0.000 | 0.003 | 0.032 | 3826 |
| GO_REGULATION_OF_ MICROTUBULE_ POLYMERIZATION_OR_ DEPOLYMERIZATION | 138 | 0.40 | 1.64 | 0.003 | 0.004 | 0.056 | 3222 |
| GO_NUCLEAR_ CHROMOSOME | 438 | 0.35 | 1.61 | 0.000 | 0.005 | 0.082 | 3830 |
| GO_CHROMOSOME_ TELOMERIC_REGION | 129 | 0.39 | 1.59 | 0.005 | 0.005 | 0.101 | 4619 |
| GO_NUCLEAR_ CHROMOSOME_ TELOMERIC_REGION | 103 | 0.36 | 1.46 | 0.017 | 0.013 | 0.336 | 4619 |
| GO_CONDENSED_NUCLEAR _CHROMOSOME | 71 | 0.35 | 1.29 | 0.108 | 0.047 | 0.872 | 3830 |
| GO_CHROMATIN | 374 | 0.35 | 1.61 | 0.000 | 0.004 | 0.010 | 3812 |
| GO_CHROMOSOME_ SEGREGATION | 213 | 0.46 | 2.01 | 0.000 | 0.000 | 0.001 | 3798 |
| GO_NUCLEAR_ CHROMOSOME_ SEGREGATION | 178 | 0.45 | 1.90 | 0.000 | 0.002 | 0.006 | 3798 |
| GO_CHROMOSOME_ ORGANIZATION | 821 | 0.40 | 1.89 | 0.000 | 0.001 | 0.007 | 4255 |
| GO_MEIOTIC_CELL_CYCLE | 152 | 0.34 | 1.44 | 0.014 | 0.033 | 0.432 | 4199 |

#### Supplementary Table 4. Organelle organisation gene sets enriched in A375M2.

| Organelle organisation | | | | | | | |
|---|---|---|---|---|---|---|---|
| **GS follow link to MSigDB** | **SIZE** | **ES** | **NES** | **NOM p-val** | **FDR q-val** | **FWER p-val** | **RANK AT MAX** |
| GO_NUCLEUS_ ORGANIZATION | 114 | 0.42 | 1.69 | 0.001 | 0.003 | 0.037 | 4158 |
| GO_ENDOMEMBRANE_ SYSTEM_ORGANIZATION | 398 | 0.36 | 1.65 | 0.000 | 0.003 | 0.052 | 4160 |
| GO_NEGATIVE_REGULATION _OF_ORGANELLE_ ORGANIZATION | 321 | 0.36 | 1.64 | 0.000 | 0.004 | 0.059 | 3881 |
| GO_NUCLEAR_ENVELOPE_ ORGANIZATION | 66 | 0.43 | 1.60 | 0.009 | 0.005 | 0.087 | 4134 |
| GO_REGULATION_OF_ ORGANELLE_ORGANIZATION | 988 | 0.32 | 1.54 | 0.000 | 0.007 | 0.162 | 3919 |
| GO_NEGATIVE_REGULATION _OF_CELLULAR_ COMPONENT_ ORGANIZATION | 593 | 0.29 | 1.38 | 0.002 | 0.024 | 0.599 | 3778 |
| GO_MEMBRANE_ ORGANIZATION | 778 | 0.27 | 1.31 | 0.003 | 0.041 | 0.829 | 4245 |

#### Supplementary Table 5. Cellular localisation gene sets enriched in A375M2.

| Cellular localisation | | | | | | | |
|---|---|---|---|---|---|---|---|
| **GS follow link to MSigDB** | **SIZE** | **ES** | **NES** | **NOM p-val** | **FDR q-val** | **FWER p-val** | **RANK AT MAX** |
| GO_PROTEIN_LOCALIZATION _TO_ORGANELLE | 490 | 0.36 | 1.66 | 0.000 | 0.003 | 0.046 | 4324 |
| GO_ORGANELLE_ LOCALIZATION | 358 | 0.33 | 1.51 | 0.000 | 0.009 | 0.216 | 4523 |
| GO_SINGLE_ORGANISM_ CELLULAR_LOCALIZATION | 798 | 0.31 | 1.46 | 0.000 | 0.012 | 0.313 | 4523 |
| GO_ESTABLISHMENT_OF_ LOCALIZATION_IN_CELL | 1449 | 0.30 | 1.43 | 0.000 | 0.016 | 0.403 | 4523 |
| GO_PROTEIN_LOCALIZATION | 1565 | 0.29 | 1.43 | 0.000 | 0.016 | 0.412 | 4250 |
| GO_PROTEIN_LOCALIZATION _TO_NUCLEUS | 142 | 0.34 | 1.40 | 0.024 | 0.019 | 0.501 | 3746 |
| GO_CELLULAR_ MACROMOLECULE_ LOCALIZATION | 1079 | 0.31 | 1.48 | 0.000 | 0.028 | 0.355 | 4335 |

#### Supplementary Table 6. Cellular development gene sets enriched in A375M2.

| Cellular development | | | | | | | |
|---|---|---|---|---|---|---|---|
| **GS follow link to MSigDB** | **SIZE** | **ES** | **NES** | **NOM p-val** | **FDR q-val** | **FWER p-val** | **RANK AT MAX** |
| GO_HEART_ MORPHOGENESIS | 197 | 0.46 | 1.96 | 0.000 | 0.000 | 0.000 | 3003 |
| GO_ANATOMICAL_STRUCTU RE_FORMATION_INVOLVED_ IN_MORPHOGENESIS | 843 | 0.39 | 1.86 | 0.000 | 0.001 | 0.002 | 3447 |
| GO_CIRCULATORY_SYSTEM _DEVELOPMENT | 717 | 0.38 | 1.77 | 0.000 | 0.004 | 0.010 | 3447 |
| GO_HEART_DEVELOPMENT | 423 | 0.38 | 1.75 | 0.000 | 0.003 | 0.012 | 3019 |
| GO_ORGAN_ MORPHOGENESIS | 780 | 0.36 | 1.72 | 0.000 | 0.004 | 0.016 | 3503 |
| GO_TISSUE_DEVELOPMENT | 1363 | 0.32 | 1.55 | 0.000 | 0.010 | 0.092 | 3344 |
| GO_MUSCLE_TISSUE_ DEVELOPMENT | 243 | 0.34 | 1.51 | 0.002 | 0.011 | 0.127 | 3015 |
| GO_MUSCLE_STRUCTURE_ DEVELOPMENT | 387 | 0.31 | 1.41 | 0.002 | 0.026 | 0.324 | 4477 |
| GO_MUSCLE_ORGAN_ DEVELOPMENT | 247 | 0.32 | 1.41 | 0.007 | 0.026 | 0.332 | 3049 |
| GO_NEUROGENESIS | 1251 | 0.29 | 1.41 | 0.000 | 0.024 | 0.335 | 3521 |
| GO_SKELETAL_MUSCLE_ CELL_DIFFERENTIATION | 49 | 0.40 | 1.37 | 0.070 | 0.027 | 0.427 | 3015 |
| GO_OSSIFICATION | 231 | 0.41 | 1.80 | 0.000 | 0.003 | 0.022 | 3335 |

#### Supplementary Table 7. Cytoskeleton organisation gene sets enriched in A375M2.

| Cytoskeleton organisation | | | | | | | |
|---|---|---|---|---|---|---|---|
| **GS follow link to MSigDB** | **SIZE** | **ES** | **NES** | **NOM p-val** | **FDR q-val** | **FWER p-val** | **RANK AT MAX** |
| GO_MICROTUBULE_CYTOSKELETON_ORGANIZATION | 277 | 0.42 | 1.87 | 0.000 | 0.001 | 0.004 | 3074 |
| GO_NEGATIVE_REGULATION_OF_CYTOSKELETON_ORGANIZATION | 182 | 0.37 | 1.58 | 0.001 | 0.005 | 0.104 | 2971 |
| GO_CYTOSKELETON_ORGANIZATION | 714 | 0.33 | 1.58 | 0.000 | 0.005 | 0.109 | 4134 |
| GO_REGULATION_OF_CYTOSKELETON_ORGANIZATION | 422 | 0.34 | 1.57 | 0.000 | 0.005 | 0.114 | 3271 |
| GO_CYTOSKELETAL_PART | 1125 | 0.33 | 1.60 | 0.000 | 0.011 | 0.048 | 3065 |
| GO_CYTOSKELETON | 1571 | 0.32 | 1.55 | 0.000 | 0.010 | 0.064 | 3074 |
| GO_CYTOSKELETAL_PROTEIN_BINDING | 727 | 0.30 | 1.42 | 0.000 | 0.022 | 0.268 | 4255 |

#### Supplementary Table 8. Cellular movement gene sets enriched in A375M2.

| Cellular movement | | | | | | | |
|---|---|---|---|---|---|---|---|
| **GS follow link to MSigDB** | **SIZE** | **ES** | **NES** | **NOM p-val** | **FDR q-val** | **FWER p-val** | **RANK AT MAX** |
| GO_REGULATION_OF_CELLULAR_COMPONENT_MOVEMENT | 710 | 0.30 | 1.42 | 0.000 | 0.016 | 0.432 | 3920 |
| GO_POSITIVE_REGULATION_OF_LOCOMOTION | 392 | 0.30 | 1.36 | 0.002 | 0.027 | 0.651 | 3920 |
| GO_MOVEMENT_OF_CELL_OR_SUBCELLULAR_COMPONENT | 1151 | 0.27 | 1.33 | 0.001 | 0.036 | 0.779 | 3845 |

#### Supplementary Table 9. Cellular transport gene sets enriched in A375M2.

| Cellular transport | | | | | | | |
|---|---|---|---|---|---|---|---|
| **GS follow link to MSigDB** | **SIZE** | **ES** | **NES** | **NOM p-val** | **FDR q-val** | **FWER p-val** | **RANK AT MAX** |
| GO_CYTOSKELETON_DEPENDENT_INTRACELLULAR_TRANSPORT | 100 | 0.50 | 2.00 | 0.000 | 0.000 | 0.001 | 3872 |

#### Supplementary Table 10. Membrane remodelling gene sets enriched in A375M2.

| Membrane remodelling | | | | | | | |
|---|---|---|---|---|---|---|---|
| **GS follow link to MSigDB** | **SIZE** | **ES** | **NES** | **NOM p-val** | **FDR q-val** | **FWER p-val** | **RANK AT MAX** |
| GO_ORGANELLE_FISSION | 389 | 0.39 | 1.81 | 0.000 | 0.003 | 0.018 | 4177 |

#### Supplementary Table 11. Nuclear matrix gene sets enriched in A375M2.

| Nuclear matrix | | | | | | | |
|---|---|---|---|---|---|---|---|
| **GS follow link to MSigDB** | **SIZE** | **ES** | **NES** | **NOM p-val** | **FDR q-val** | **FWER p-val** | **RANK AT MAX** |
| GO_NUCLEAR_MATRIX | 89 | 0.55 | 2.15 | 0.000 | 0.000 | 0.000 | 3756 |

#### Supplementary Table 12. Biosynthesis gene sets enriched in A375M2.

| Biosynthesis | | | | | | | |
|---|---|---|---|---|---|---|---|
| **GS follow link to MSigDB** | **SIZE** | **ES** | **NES** | **NOM p-val** | **FDR q-val** | **FWER p-val** | **RANK AT MAX** |
| GO_STEROL_BIOSYNTHETIC_PROCESS | 41 | 0.55 | 1.85 | 0.000 | 0.005 | 0.005 | 1538 |
| GO_ALCOHOL_BIOSYNTHETIC_PROCESS | 97 | 0.38 | 1.47 | 0.022 | 0.023 | 0.191 | 5032 |

#### Supplementary Table 13. Clinical information for primary melanoma patients.

| | Cohort A patients | Cohort B patients |
|---|---|---|
| **Age, years** (mean ± SD) | 67 ± 16.5 | 61 ± 18 |
| **Gender** | | |
| Male | 10 (52.5%) | 15 (51.7%) |
| Female | 9 (47.4%) | 14 (48.3%) |
| **Location** | | |
| Trunk | 7 (36.8%) | 16 (55.2%) |
| Head and neck | 5 (26.4%) | 7 (24.2%) |
| Lower limb | 3 (15.8%) | 2 (6.9%) |
| Upper limb | 2 (10.5%) | 1 (3.4%) |
| Foot | 2 (10.5%) | 2 (6.9%) |
| Hand | | 1 (3.4%) |

#### Supplementary Table 14. Clinical information for metastatic melanoma patients.

| | Cohort A patients | Cohort B patients |
|---|---|---|
| **Age, years** (mean ± SD) | 67 ± 13 | 61 ± 18 |
| **Gender** | | |
| Male | 9 (64.3%) | 15 (51.7%) |
| Female | 5 (35.7%) | 14 (48.3%) |
| **Location** | | |
| Lymph node | 9 (64.3%) | 29 (100%) |
| Cutaneous/subcutaneous | 4 (28.6%) | |
| Lung | 1 (7.1%) | |

#### Supplementary Table 15. List of PCR primer sequences.

| PCR primer | PCR primer sequence |
|---|---|
| LAP1B FWD | 5’-ATATGAATTCGCCACCATGGCGGG-3’ |
| LAP1B REV | 5’-TATGCGGCCGCTTAAGCAGATGC-3’ |
| LAP1C FWD | 5’-ATATGAATTCGCCACCATGAAGACGCGAAGGACTACC-3’ |
| LAP1B M122A FWD | 5’-GGAAACCGAGGAAGCGAAGACGCGAAGG-3’ |
| LAP1B M122A REV | 5’-CCTTCGCGTCTTCGCTTCCTCGGTTTCC-3’ |
| LAP1B Δ1-72 FWD | 5’- ATATGAATTCGCCACCATGCTGGTGGCCAAAGAAAGG-3' |
| LAP1 siRNA#1 R FWD | 5’-GCGAAAGAGGAAGTAAGGGAGAGTGCATATTACCTTCGGTCTAGG-3’ |
| LAP1 siRNA#1 R REV | 5’-GCCTAGACCGAAGGTAATATGCACTCTCCCTTACTTCCTCTTTCG-3’ |

#### Supplementary Table 16. List of siRNA sequences.

| Target | siRNA sequence 5’ to 3’ |
|---|---|
| Control siRNA | Non-targeting siRNA: D-001810-01-20 |
| LAP1 | siGENOME SMARTpool:<br>5’-GAACUGAGCAAUGGAUUUA-3’<br>5’-UAAGAUACCCAAGAUAUGA-3’<br>5’-GAAGAGUUCCGGUCCGAUU-3’<br>5’-UGAGAGAAAGCGCGUACUA-3’ |
| | ON-TARGETplus individual sequence:<br>5’-UGAGAGAAAGCGCGUACUA-3’<br>5’-GGUCCGAUUCUGCGAAAGA-3’<br>5’-GAGGAAAUGAAGACGCGAA-3’<br>5’-CGUCUUUCCUCUAGUACUA-3’ |
| LAMIN A/C | siGENOME SMARTpool:<br>5’-GAAGGAGGGUGACCUGAUA-3’<br>5’-GAGCUGAAAGCGCGCAAUA-3’<br>5’-GUACGGCUCUCAUCAACUC-3’<br>5’-CUGGGCAGGUGGUGACGAU-3’ |
| LAMIN B1 | siGENOME SMARTpool:<br>5’-GAAGGAAUCUGAUCUUAAU-3’<br>5’-CAACUGACCUCAUCUGGAA-3’<br>5’-GAAAGAGUCUAGAGCAUGU-3’<br>5’-GCAUGAAACGCGCUUGGUA-3’ |
| LAMIN B2 | siGENOME SMARTpool:<br>5’-GGAGAUCGCCUACAAGUUC-3’<br>5’-GAACAACUCGGACAAGGAU-3’<br>5’-UAACGCGGAUGGCGAGGAA-3’<br>5’-CCUCGACGCUGGUGUGGAA-3’ |

### Supplementary Table 17. List of BioWave progam details.

| Description | Step# | Time (min) | Time (sec) | Power (Watts) | SteadyTemp temperature (ºC) | Vacuum cycle vent time (sec) | Vacuum cycle vacuum time (sec) | Vacuum set point (inch Hg) | User Prompt (1 = YES, 0 = NO) | Vacuum OFF (1 = no vacuum, 0 = vacuum) | Vacuum cycle (1 = ON, 0 = OFF) | Vacuum ON (1 = vacuum, 0 = no vacuum) |
|---|---|---|---|---|---|---|---|---|---|---|---|---|
| BENCH STEP Rinse in 0.1M PB | 1 | 0 | 0 | 0 | 21 | 0 | 0 | 0 | 1 | 1 | 0 | 0 |
| BENCH STEP Rinse in 0.1M PB | 2 | 0 | 0 | 0 | 21 | 0 | 0 | 0 | 1 | 1 | 0 | 0 |
| Rinse in 0.1M PB | 3 | 0 | 40 | 250 | 21 | 0 | 0 | 0 | 1 | 1 | 0 | 0 |
| Rinse in 0.1M PB | 4 | 0 | 40 | 250 | 21 | 0 | 0 | 0 | 1 | 1 | 0 | 0 |
| Osmium ON | 5 | 2 | 0 | 100 | 21 | 0 | 0 | 20 | 1 | 0 | 0 | 1 |
| Osmium OFF | 6 | 2 | 0 | 0 | 21 | 0 | 0 | 20 | 0 | 0 | 0 | 1 |
| Osmium ON | 7 | 2 | 0 | 100 | 21 | 0 | 0 | 20 | 0 | 0 | 0 | 1 |
| Osmium OFF | 8 | 2 | 0 | 0 | 21 | 0 | 0 | 20 | 0 | 0 | 0 | 1 |
| Osmium ON | 9 | 2 | 0 | 100 | 21 | 0 | 0 | 20 | 0 | 0 | 0 | 1 |
| Osmium OFF | 10 | 2 | 0 | 0 | 21 | 0 | 0 | 20 | 0 | 0 | 0 | 1 |
| Osmium ON | 11 | 2 | 0 | 100 | 21 | 0 | 0 | 20 | 0 | 0 | 0 | 1 |
| BENCH STEP Rinse in water | 12 | 0 | 0 | 0 | 21 | 0 | 0 | 0 | 1 | 1 | 0 | 0 |
| BENCH STEP Rinse in water | 13 | 0 | 0 | 0 | 21 | 0 | 0 | 0 | 1 | 1 | 0 | 0 |
| Rinse in water | 14 | 0 | 40 | 250 | 21 | 0 | 0 | 0 | 1 | 1 | 0 | 0 |
| Rinse in water | 15 | 0 | 40 | 250 | 21 | 0 | 0 | 0 | 1 | 1 | 0 | 0 |
| Thiocarbohydrazide ON | 16 | 2 | 0 | 100 | 40 | 0 | 0 | 20 | 1 | 0 | 0 | 1 |
| Thiocarbohydrazide OFF | 17 | 2 | 0 | 0 | 40 | 0 | 0 | 20 | 0 | 0 | 0 | 1 |
| Thiocarbohydrazide ON | 18 | 2 | 0 | 100 | 40 | 0 | 0 | 20 | 0 | 0 | 0 | 1 |
| Thiocarbohydrazide OFF | 19 | 2 | 0 | 0 | 40 | 0 | 0 | 20 | 0 | 0 | 0 | 1 |
| Thiocarbohydrazide ON | 20 | 2 | 0 | 100 | 40 | 0 | 0 | 20 | 0 | 0 | 0 | 1 |
| Thiocarbohydrazide OFF | 21 | 2 | 0 | 0 | 40 | 0 | 0 | 20 | 0 | 0 | 0 | 1 |
| Thiocarbohydrazide ON | 22 | 2 | 0 | 100 | 40 | 0 | 0 | 20 | 0 | 0 | 0 | 1 |
| BENCH STEP Rinse in water | 23 | 0 | 0 | 0 | 21 | 0 | 0 | 0 | 1 | 1 | 0 | 0 |
| BENCH STEP Rinse in water | 24 | 0 | 0 | 0 | 21 | 0 | 0 | 0 | 1 | 1 | 0 | 0 |
| Rinse in water | 25 | 0 | 40 | 250 | 21 | 0 | 0 | 0 | 1 | 1 | 0 | 0 |
| Rinse in water | 26 | 0 | 40 | 250 | 21 | 0 | 0 | 0 | 1 | 1 | 0 | 0 |
| Osmium ON | 27 | 2 | 0 | 100 | 21 | 0 | 0 | 20 | 1 | 0 | 0 | 1 |
| Osmium OFF | 28 | 2 | 0 | 0 | 21 | 0 | 0 | 20 | 0 | 0 | 0 | 1 |
| Osmium ON | 29 | 2 | 0 | 100 | 21 | 0 | 0 | 20 | 0 | 0 | 0 | 1 |
| Osmium OFF | 30 | 2 | 0 | 0 | 21 | 0 | 0 | 20 | 0 | 0 | 0 | 1 |
| Osmium ON | 31 | 2 | 0 | 100 | 21 | 0 | 0 | 20 | 0 | 0 | 0 | 1 |
| Osmium OFF | 32 | 2 | 0 | 0 | 21 | 0 | 0 | 20 | 0 | 0 | 0 | 1 |
| Osmium ON | 33 | 2 | 0 | 100 | 21 | 0 | 0 | 20 | 0 | 0 | 0 | 1 |
| BENCH STEP Rinse in water | 34 | 0 | 0 | 0 | 21 | 0 | 0 | 0 | 1 | 1 | 0 | 0 |
| BENCH STEP Rinse in water | 35 | 0 | 0 | 0 | 21 | 0 | 0 | 0 | 1 | 1 | 0 | 0 |
| Rinse in water | 36 | 0 | 40 | 250 | 21 | 0 | 0 | 0 | 1 | 1 | 0 | 0 |
| Rinse in water | 37 | 0 | 40 | 250 | 21 | 0 | 0 | 0 | 1 | 1 | 0 | 0 |
| Uranyl acetate ON | 38 | 2 | 0 | 100 | 40 | 0 | 0 | 20 | 1 | 0 | 0 | 1 |
| Uranyl acetate OFF | 39 | 2 | 0 | 0 | 40 | 0 | 0 | 20 | 0 | 0 | 0 | 1 |
| Uranyl acetate ON | 40 | 2 | 0 | 100 | 40 | 0 | 0 | 20 | 0 | 0 | 0 | 1 |
| Uranyl acetate OFF | 41 | 2 | 0 | 0 | 40 | 0 | 0 | 20 | 0 | 0 | 0 | 1 |
| Uranlyl acetate ON | 42 | 2 | 0 | 100 | 40 | 0 | 0 | 20 | 0 | 0 | 0 | 1 |
| Uranyl acetate OFF | 43 | 2 | 0 | 0 | 40 | 0 | 0 | 20 | 0 | 0 | 0 | 1 |
| Uranyl acetate ON | 44 | 2 | 0 | 100 | 40 | 0 | 0 | 20 | 0 | 0 | 0 | 1 |
| BENCH STEP Rinse in water | 45 | 0 | 0 | 0 | 40 | 0 | 0 | 0 | 1 | 1 | 0 | 0 |
| BENCH STEP Rinse in water | 46 | 0 | 0 | 0 | 40 | 0 | 0 | 0 | 1 | 1 | 0 | 0 |
| Rinse in water | 47 | 0 | 45 | 250 | 40 | 0 | 0 | 0 | 1 | 1 | 0 | 0 |
| Rinse in water | 48 | 0 | 45 | 250 | 40 | 0 | 0 | 0 | 1 | 1 | 0 | 0 |
| Lead aspartate ON | 49 | 2 | 0 | 100 | 50 | 0 | 0 | 20 | 1 | 0 | 0 | 1 |
| Lead aspartate OFF | 50 | 2 | 0 | 0 | 50 | 0 | 0 | 20 | 0 | 0 | 0 | 1 |
| Lead aspartate ON | 51 | 2 | 0 | 100 | 50 | 0 | 0 | 20 | 0 | 0 | 0 | 1 |
| Lead aspartate OFF | 52 | 2 | 0 | 0 | 50 | 0 | 0 | 20 | 0 | 0 | 0 | 1 |
| Lead aspartate ON | 53 | 2 | 0 | 100 | 50 | 0 | 0 | 20 | 0 | 0 | 0 | 1 |
| Lead aspartate OFF | 54 | 2 | 0 | 0 | 50 | 0 | 0 | 20 | 0 | 0 | 0 | 1 |
| Lead aspartate ON | 55 | 2 | 0 | 100 | 50 | 0 | 0 | 20 | 0 | 0 | 0 | 1 |
| BENCH STEP Rinse in water | 56 | 0 | 0 | 0 | 21 | 0 | 0 | 0 | 1 | 1 | 0 | 0 |
| BENCH STEP Rinse in water | 57 | 0 | 0 | 0 | 21 | 0 | 0 | 0 | 1 | 1 | 0 | 0 |
| Rinse in water | 58 | 0 | 45 | 250 | 21 | 0 | 0 | 0 | 1 | 1 | 0 | 0 |
| Rinse in water | 59 | 0 | 45 | 250 | 21 | 0 | 0 | 0 | 1 | 1 | 0 | 0 |
| 70% Ethanol ON | 60 | 0 | 40 | 250 | 21 | 0 | 0 | 0 | 1 | 1 | 0 | 0 |
| 70% Ethanol ON | 61 | 0 | 40 | 250 | 21 | 0 | 0 | 0 | 1 | 1 | 0 | 0 |
| 90% Ethanol ON | 62 | 0 | 40 | 250 | 21 | 0 | 0 | 0 | 1 | 1 | 0 | 0 |
| 90% Ethanol ON | 63 | 0 | 40 | 250 | 21 | 0 | 0 | 0 | 1 | 1 | 0 | 0 |
| 100% Ethanol ON | 64 | 0 | 40 | 250 | 21 | 0 | 0 | 0 | 1 | 1 | 0 | 0 |
| 100% Ethanol ON | 65 | 0 | 40 | 250 | 21 | 0 | 0 | 0 | 1 | 1 | 0 | 0 |
| 50% Resin ON | 66 | 3 | 0 | 250 | 21 | 30 | 30 | 20 | 1 | 0 | 1 | 0 |
| 100% Resin ON | 67 | 3 | 0 | 250 | 21 | 30 | 30 | 20 | 1 | 0 | 1 | 0 |
| 100% Resin ON | 68 | 3 | 0 | 250 | 21 | 30 | 30 | 20 | 1 | 0 | 1 | 0 |
| 100% Resin ON | 69 | 3 | 0 | 250 | 21 | 30 | 30 | 20 | 1 | 0 | 1 | 0 |
| 100% Resin ON | 70 | 3 | 0 | 250 | 21 | 30 | 30 | 20 | 1 | 0 | 1 | 0 |
| TURN SYSTEM OFF | 71 | 0 | 0 | 0 | 21 | 0 | 0 | 0 | 0 | 1 | 0 | 0 |
